# Seed dispersal disruption limits tropical forest regrowth

**DOI:** 10.1101/2024.12.06.627256

**Authors:** Evan C. Fricke, Susan C. Cook-Patton, Charles F. Harvey, César Terrer

## Abstract

Many trees depend on animals for seed dispersal, and human activities that disrupt seed dispersal by animals may impact forest regeneration and carbon storage. Yet whether expected negative impacts are observable across regrowing forests remains untested. We modeled seed dispersal disruption and its relationship to aboveground carbon accumulation observed across 3026 sites in the tropics, where most trees are animal dispersed. We found that seed dispersal disruption explains wide variation in local carbon accumulation rates. Across areas identified for restoration, we estimate that seed dispersal disruption reduces carbon accumulation rates by 57% on average. These results advance understanding of animal biodiversity’s impact on forest carbon and emphasize the need to address biodiversity loss and climate change together.

---

Climate change and biodiversity loss are critical environmental challenges of our time. Well-established linkages between these challenges include the exacerbating effect of climate change on biodiversity loss (*1*) and the positive relationship between plant biodiversity and carbon storage (*2*, *3*) in vegetation communities. Less well known is the potential for terrestrial vertebrates such as birds and mammals to influence carbon storage at a scale that is relevant to climate change (*4–7*), yet animals may influence the carbon stored in forest ecosystems through their role as seed dispersers.

Many tree species rely on animals for the dispersal of their seeds (*8*). Without dispersal, plants have lower germination, growth, and survival (*9*, *10*), and they often cannot access suitable regeneration sites including those made available through disturbances such as fire, storms, drought, and deforestation (*11*, *12*). In forests regrowing after disturbance, dispersal limitations pose a primary barrier to regeneration (*13*). Where seed dispersers have declined, field studies show reduced abundance and diversity of regenerating trees (*14*) and altered functional composition, especially the lower prevalence of late successional, large-seeded trees with dense wood (*15–17*). Together, local ecological research shows that seed dispersers affect the abundance, diversity, and functional composition of regenerating trees.

The disruption of seed dispersal—caused by ongoing declines in animal diversity, abundance, and movement (*18–20*)—could therefore negatively impact forest carbon dynamics. Process-based models, which have paired short-term field data with long-term demographic modelling, suggest that seed dispersal disruption should reduce carbon storage in both existing forests (*21*, *22*) and areas regrowing after disturbance (*23*). Although these studies have been restricted to local areas, they indicate that the magnitude of seed dispersal disruption’s negative effects may be large, particularly in regrowing forests. For example, in a fragmented region of the Atlantic Forest in Brazil, modelling by Bello *et al.* (*23*) showed that observed disruption of seed dispersal would lead to 38% lower carbon storage in naturally regrowing forests in the more fragmented versus less fragmented areas of the study landscape. Yet whether the signature of seed dispersal disruption that is predicted by such models is discernible across field records of tropical forest regrowth globally remains untested. This obscures the relevance of animal seed dispersers for broad-scale carbon dynamics, especially the impact of seed dispersal disruption on natural forest regrowth, which contributes substantially to the land carbon sink (*24*, *25*) and which is often promoted as a natural climate solution (*26*) to be implemented alongside necessary steep reductions in fossil fuel emissions.

## Dependence on animal seed dispersal

To assess how seed dispersal disruption may affect natural forest regrowth, we first asked how prevalent animal-mediated seed dispersal is across all forest and savanna biomes. We developed a relative metric of the prevalence of animal-mediated seed dispersal that is weighted by plant species’ abundance and their dependence on animals for seed dispersal (*27*). We analyzed data from 17,071 vegetation plots on the relative abundance of tree species and their dependence on animals for seed dispersal. Species-specific estimates of dependence on animals for seed dispersal were developed using 346,501 records of seed dispersal mode for 24,600 plant species from 4,693 genera and 369 families. The prevalence of seed dispersal by animals varies spatially (*R*^2^ = 0.66 on test data; map in Fig. 1; Fig. S1) and strongly increases toward the equator (*F* = 1893, *P* < 0.0001; inset plot in Fig. 1). On average across tropical plots, we estimate that 81% of trees rely on animals for seed dispersal. This suggests that loss of animal seed dispersal function could limit the regrowth and carbon accumulation potential of tropical forests in particular. We therefore focus our next analyses on tropical regions where animal seed dispersers may be most influential for natural regrowth.

**Fig. 1.**
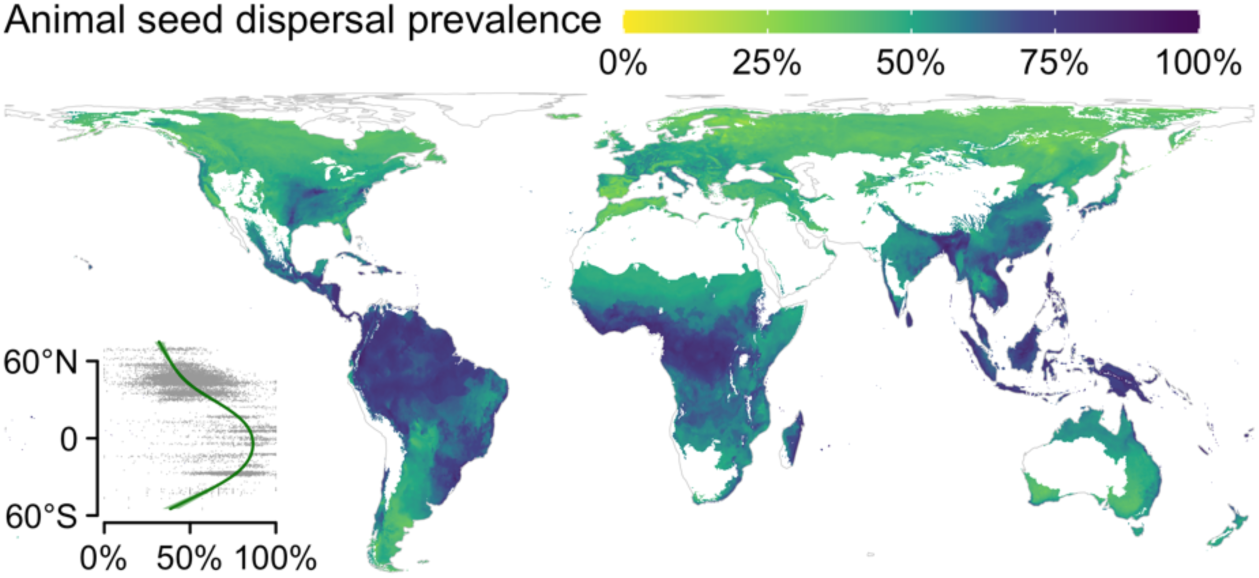
The prevalence of seed dispersal by animals in woody vegetation across forest and savanna biomes. Points in the inset panel show the weighted, relative metric of animal seed dispersal prevalence in 17071 vegetation plots across latitude and the fitted line shows the relationship with latitude from a generalized additive model. The map illustrates estimated animal dispersal prevalence developed using the plot data, climate covariates, and a random forest model. Estimates of model uncertainty are presented in Fig. S1.

## Seed dispersal disruption

A key challenge for evaluating how seed dispersal disruption may influence tropical forest regrowth is the limited understanding of how seed dispersal function varies across landscapes; whereas global gridded products of environmental variables (e.g., temperature, precipitation) are available for modelling forest processes (*25*, *26*), analogous estimates for seed dispersal disruption are lacking. To generate location-specific estimates, we combined published estimates of animal species-specific seed dispersal function, compilations of field data on animal presence and movement, and geographic distribution data for mammals and birds (*27*) (see conceptual diagram showing an overview of these methods in Fig. S2). We started with published estimates of the seed dispersal function provided by birds and mammals globally from ref. (*28*). Their modelling involved ecological analyses of the quantity of seeds that animals disperse (406 local inventories of seed dispersal interactions), the distance they carry seeds (data on seed retention times and seed dispersal distance for interactions involving 302 animal species), and the probability that seeds germinate following deposition (results of 2215 studies of the impact of gut passage on seed germination). Their modelling showed a high capacity to predict aspects of seed dispersal based on species’ traits (*28*), such as seed retention times (cross-validation *R*^2^ = 0.96; Fig. 2A). Together, this modelling offers species-specific estimates of seed dispersal function or, across a local species assemblage, the seed dispersal function provided by birds and mammals at a location. However, these estimates do not account for anthropogenic impacts on seed dispersers’ presence or movement patterns. To incorporate these impacts, here we analyzed the effect of habitat fragmentation on species presence (*27*) (61,716 occurrence records across 1990 species; Fig. S3A) and, following ref. (*19*), of the magnitude of the human footprint on animal movement (488,583 geolocations across 1838 individuals of 80 species; Fig. S3B). This showed, for example, that when the human footprint index is higher, animals move shorter distances during the time when they retain seeds (Fig. 2B). Using data on the geographic ranges of birds and mammals, habitat fragmentation, and human footprint, we then estimated seed dispersal disruption as the proportional loss of seed dispersal function attributable to these anthropogenic drivers across tropical regions (*27*) (Fig. S2). This metric emphasizes the quantity of effective seed dispersal, which is important for overcoming the seed limitation that often hampers regrowth (*13*), and relatively long-distance seed movements, which are important for vegetation recovery following disturbance (*11*, *29*).

**Fig. 2.**
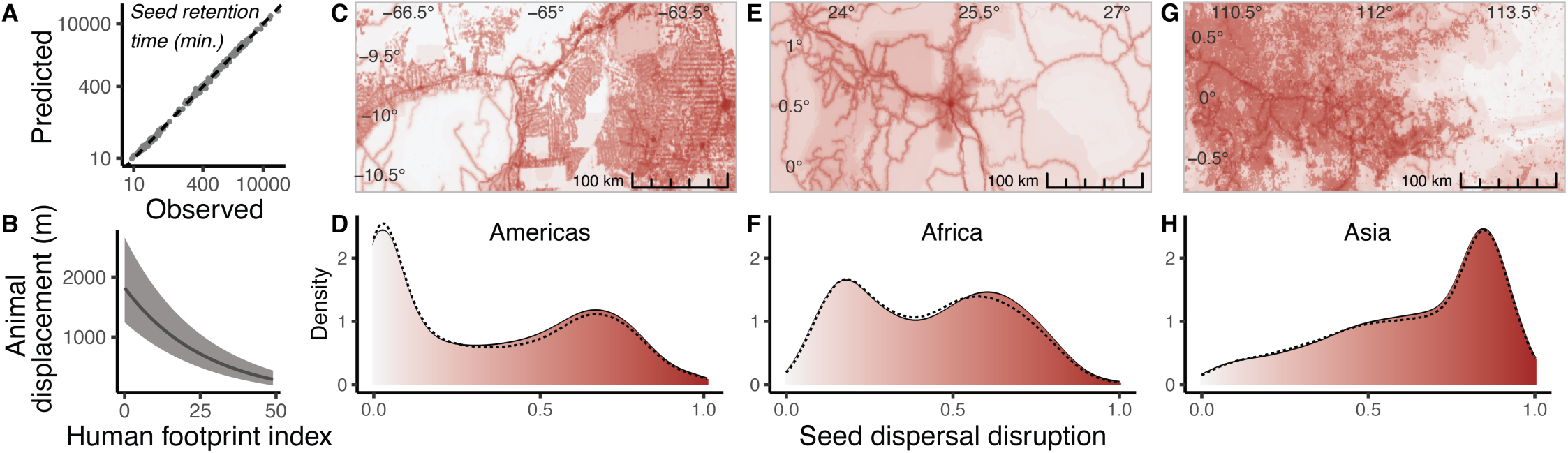
Modelling seed dispersal disruption. (**A**) Example of the processes modelled to estimate seed dispersal function, showing observed seed retention times versus model-predicted values based on species traits by ref. (*28*) (see also Fig. S2). (**A**) Observed relationship between the human footprint index and displacement distances over a typical seed retention period, based on field observations of animal movement; detailed results are presented in Figure S3. (**C-H**), examples of seed dispersal disruption values and distribution of values across regional scales; dashed lines show estimates *c.* 2000 and filled area shows estimates *c.* 2020. Inset plots show areas in the Amazon Basin (**C**), the Congo Basin (**E**), and Borneo (**G**).

**Fig. 3.**
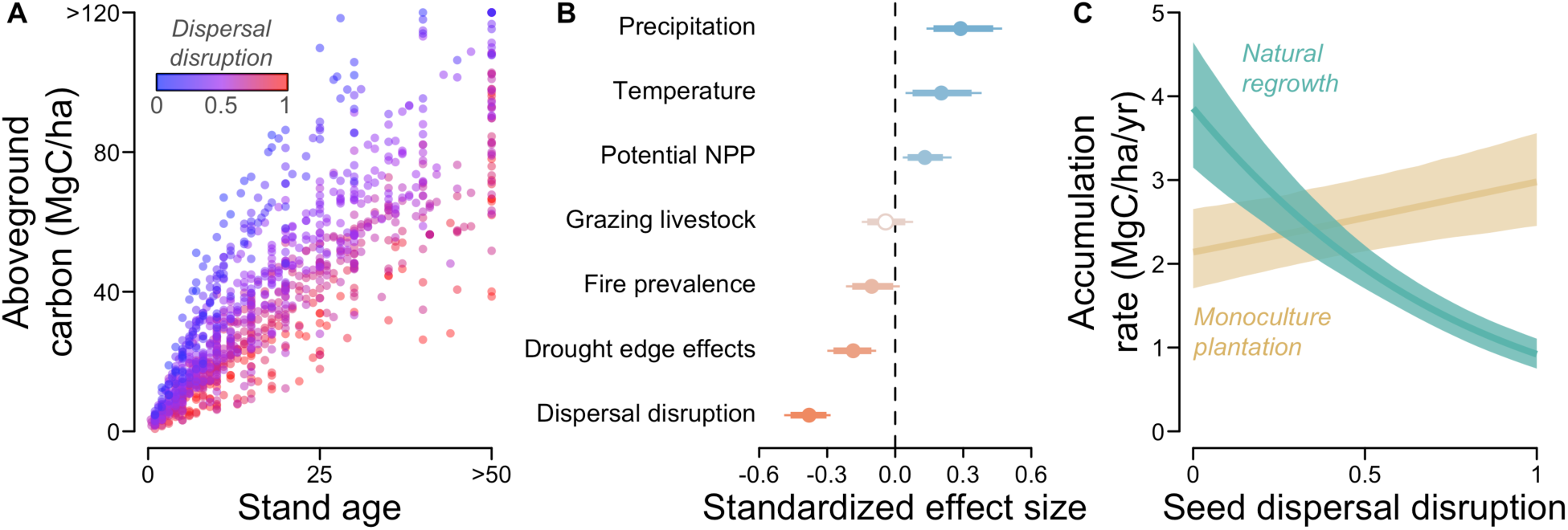
Effect of seed dispersal disruption on aboveground carbon accumulation. (**A**) Stand age versus fitted values of aboveground carbon during natural regrowth, with color gradient showing levels of seed dispersal disruption at each site. (**B**) Standardized effect sizes representing the effect of a 1 SD increase in the predictor variable on carbon accumulation rates annualized over the first 30 years of regrowth. Bars show 95% and 99% CIs, points show median posterior values, and filled points indicate 95% CIs that do not overlap zero. (**C**) Annualized carbon accumulation rate under either natural regrowth or monoculture plantation, with lines showing model estimates for average environmental conditions and shaded areas showing 95% credible intervals.

Seed dispersal disruption varies at landscape scales and across regions within tropical and subtropical forest biomes (Fig. 2C-H); key regional differences reflect factors such as the greater prevalence of high-integrity forest landscapes remaining in Amazonia and effects of widespread land conversion in Asian forest biomes (*30*, *31*). Seed dispersal disruption also changes over time under ongoing fragmentation; these changes have caused net increases in the severity of seed dispersal disruption from 2000 to 2020 (Fig. 2D,F,H).

## Impacts on carbon accumulation

We then analyzed the relationship between seed dispersal disruption and records of carbon accumulation over time from a database of field observations of natural forest regrowth in previously deforested areas (*26*). Analyzing the records of aboveground carbon in tropical plots, we fit a Bayesian model to estimate how forest carbon accumulation over time at these plots depends on seed dispersal disruption and other environmental drivers of variation in regrowth. These include processes that may also constrain regrowth in anthropogenically modified landscapes: drought conditions associated with heat- and desiccation-related edge effects, the prevalence of fire on the landscape, and the abundance of grazing livestock (*27*).

The analysis showed a strong negative effect of seed dispersal disruption on carbon accumulation of regrowing forests (Fig. 2A, Table S1; posterior estimate of effect on annualized growth: −0.38, 95% CI: [−0.46, −0.30]) and strong effects of other environmental variables (Fig. 2B; Table S1). Drought had a strong negative impact on carbon accumulation (Fig. 2B; −0.19 [−0.27, −0.10]), as did the prevalence of fire on the landscape (−0.10 [−0.19, −0.01]). A weak negative effect of grazers was not statistically significant (−0.042 [−0.123, 0.044]).

Isolating the effect of seed dispersal disruption, the fitted model estimates that areas of natural regrowth with lowest seed dispersal disruption accumulate four times as much carbon as areas with the most extreme seed dispersal disruption (Fig. 2C). This negative relationship is absent in monoculture plantation sites (*32*), where trees were planted by people (Fig. 2B, Table S1; 0.026 [0.011, 0.050]). The finding that carbon accumulation is reduced in naturally regrowing forests under more severe seed dispersal disruption is consistent with local short-term field studies that show that dispersal disruptions reduce the abundance and diversity of regrowing trees and alter their functional composition (*12–14*, *16*, *17*). Further, the observed effect on forest carbon is quantitatively consistent with the process-based modelling predictions of ref. (*23*), which were made over a moderate range in the severity of seeds dispersal disruption. Together, our results provide empirical support for large-magnitude effects of seed dispersal disruption on observed carbon accumulation across regrowing tropical forests.

## Assessing impacts on regrowth potential

Evidence for a feedback between seed dispersal disruption and forest regrowth alters our understanding of natural regrowth potential across the tropics. We highlight three factors not captured when mapping regrowth potential using only other environmental predictors (*26*, *33*). First, overlooking the impact of seed dispersal disruption may overestimate natural regrowth potential in many areas and underestimate it in others. We used our model incorporating seed dispersal disruption and environmental predictors to estimate current (*c.* 2020) regrowth potential (Fig. 4A). Compared to a model ignoring seed dispersal and including only the other environmental predictors (Fig. S4A), our model projected lower carbon accumulation in areas with greater seed dispersal disruption and, conversely, greater regrowth potential in more intact forest landscapes (Fig. S4B). Incorporating knowledge of the severity of seed dispersal can improve predictions for the carbon accumulation potential of regrowth and help to identify where natural regrowth can be most successful.

**Fig. 4.**
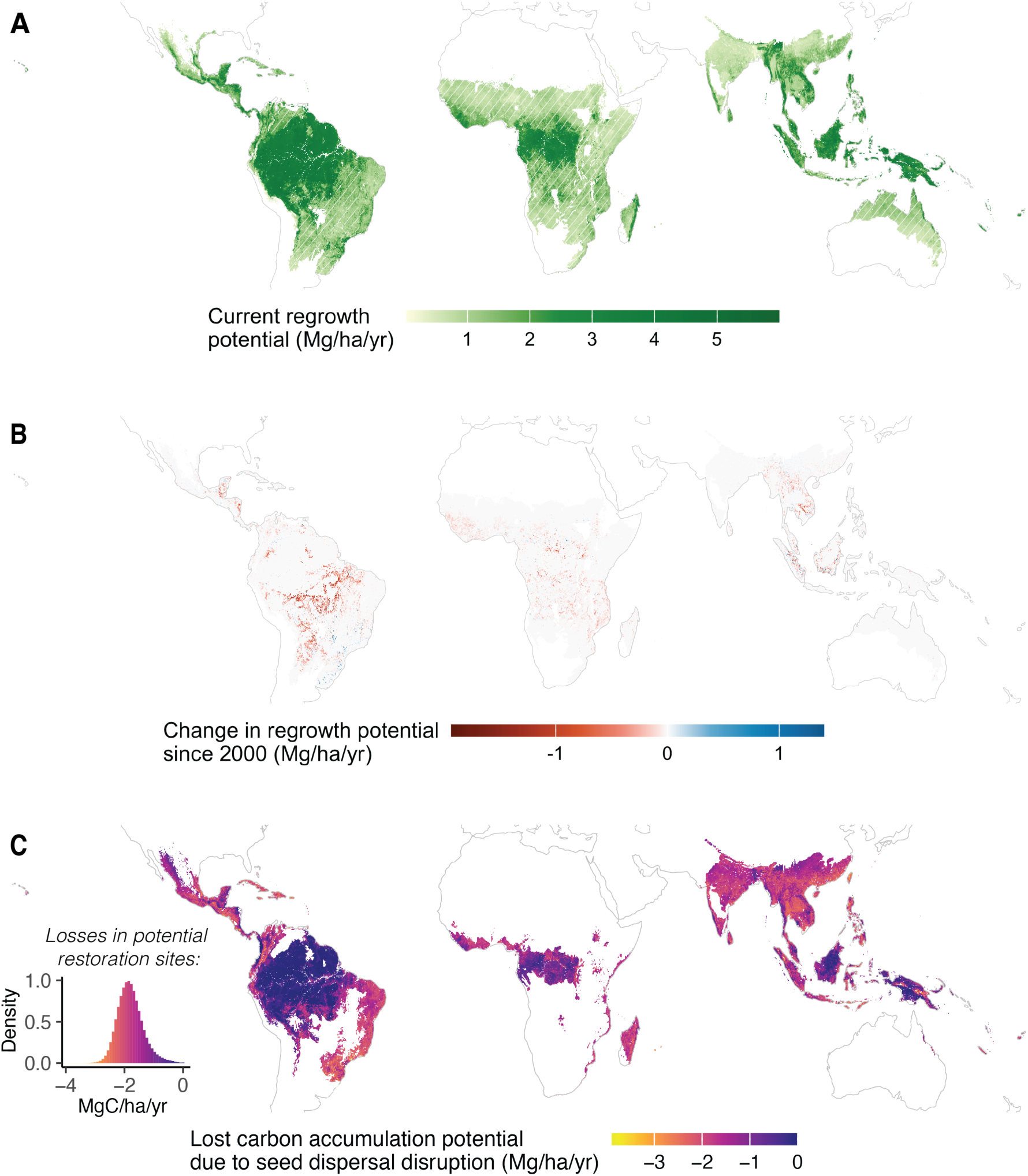
Mapping aboveground carbon accumulation potential in light of seed dispersal disruption. (**A**) The current (c. 2020) regrowth potential of tropical forest and savanna ecosystems while accounting for seed dispersal disruption. Hatching indicates savanna biomes. Estimates of model uncertainty are presented in Fig. S5. (**B**) The cumulative effect of land use change over the period 2000-2020 on natural regrowth potential. (**C**) the impact of seed dispersal disruption on regrowth potential in tropical forest areas. Values estimate the degree to which current levels of seed dispersal disruption would limit carbon accumulation for regrowing forests at each location. Inset panel shows the distribution of lost carbon accumulation potential across sites identified as potential restoration areas by ref. (*36*) that occur in the tropical forest biomes. Model uncertainty is presented in Fig. S6.

Second, because the severity of seed dispersal disruption can change over time, ongoing negative impacts on animal biodiversity could be altering the ability of tropical landscape to recover carbon through natural regrowth. For example, deforestation and reforestation both influence forest habitat fragmentation and thus the severity of seed dispersal disruption. By comparing regrowth potential for 2000 versus 2020, we estimated that tree cover changes have largely increased seed dispersal disruption and in turn reduced carbon accumulation potential (Fig. 4B). Of the areas with appreciable (over ± 5%) changes in regrowth potential, the majority (94%) showed declines. The estimated losses are conservative because many locations showing tree cover gains are plantations (*34*) that may not in reality support animal populations or movement (*35*). This indicates that ongoing pressures on animal biodiversity are reducing the ability for tropical forests to recover naturally from deforestation.

Third, we sought to understand the cumulative degree to which seed dispersal disruption has reduced the regrowth potential of tropical forest landscapes. Focusing on tropical forest biomes, we estimated regrowth potential if seed dispersal was not disrupted and compared this to current regrowth potential (Fig. 4C). In both cases, we kept other environmental variables (e.g., drought, fire) constant to isolate the effect attributable to seed dispersal disruption alone. Many areas show multiple tons per hectare of lost yearly carbon accumulation potential. To evaluate how this impacts carbon accumulation specifically in areas where forest regrowth may be implemented for ecosystem restoration and climate mitigation, we examined lost carbon accumulation potential in areas identified as potential restoration sites by ref. (*36*). We used the location of potential restoration sites not to indicate where reforestation should be implemented, but instead to characterize the impact of seed dispersal disruption in the types of landscapes where natural forest regrowth may be implemented. We estimate losses in yearly carbon accumulation averaging −1.8 Mg/ha/year (Fig. 4C inset), equating to a 57% reduction in regrowth potential for a typical site. Although these estimates represent a maximum loss—relative to the lowest levels of observed seed dispersal disruption—this result indicates that many potential restoration sites exist in landscapes where the current level of seed dispersal disruption strongly limits the climate mitigation potential of natural regrowth.

Together, these finding underscore actionable insights. First, natural regrowth projects are more likely to achieve high carbon storage if located in landscapes with low seed dispersal disruption. Target landscapes may include recently deforested areas, locations near high-integrity forest landscapes, or areas with higher existing tree cover (*37*). These are the landscapes where natural regrowth without further active restoration interventions could represent the most cost-effective restoration approach (*38*). Second, animal biodiversity decline threatens further losses in the ability of tropical forests to regrow. The protection of animal biodiversity and landscape connectivity could support the regrowth potential of tropical landscapes. A host of tools can support seed dispersal function, including habitat corridors (*39*), protected area planning that emphasizes landscape connectivity (*40*), reintroduction of seed-dispersing species (*41*), and mitigation of physical barriers to movement (*42*). Active restoration practices, such as planting tree species that attract seed dispersers to nucleate regrowth (*43*, *44*), can also accelerate recovery through the restoration of seed dispersal.

## Evaluation of our results

By modelling a key ecosystem function provided by animals and the severity of its disruption, we quantified a mechanism by which animal biodiversity changes are altering carbon dynamics across the tropics. We found that seed dispersal disruption explains four-fold variation in carbon accumulation rates across natural regrowth sites and that its current severity has reduced by more than half the average carbon accumulation potential of proposed tropical reforestation sites. Tropical regrowth forests are currently the largest land-based carbon sink (*24*) and seed disperser declines, if left unmitigated, threaten their climate mitigation potential. Our approach, and the large magnitude of the observed effects, demonstrates the potential for incorporating the ecological roles of animals within global-scale climate research and for considering the role that animal biodiversity plays in supporting forest restoration for climate mitigation. However, several limitations apply to our work that, if addressed, would advance these efforts. First, the breadth of available data on animal biodiversity has a direct impact on the precision of our conclusions. As animal biodiversity monitoring continues to improve, uncertainty and bias caused by spatiotemporal mismatches between data available on animal biodiversity and vegetation dynamics should decline. Resolving further interspecific variation in seed dispersers’ functional roles and responses to anthropogenic drivers would also offer more precise estimates of change in seed dispersal function. Second, while we were able to evaluate the effect of key drivers of forest growth, additional anthropogenic and environmental drivers could be considered in future analyses to enhance our understanding of this biodiversity-carbon linkage. Expanded data collection may help resolve the effects of differences in competing vegetation (*45*) and in other dimensions of ecosystem degradation related to differences in land use history, which did not predict natural growth potential in the data analyzed here (*26*). Other ecological functions that wild animals provide also change under the same drivers that cause seed dispersal disruption, although known impacts on processes such as pollination and herbivory under these drivers (*46*, *47*) suggest that the results we attribute to seed dispersal disruption may be conservative. Third, our analysis aims to assess key drivers of spatial variation, but the field measurements nonetheless derive from a sample of tropical areas. Research to monitor natural forest regrowth across more regions and to capture biogeographic differences in the sensitivity of forest regrowth to animal seed dispersal disruption could reduce spatial biases.

Our study demonstrates a strong linkage between animal biodiversity and terrestrial carbon storage and identifies animal biodiversity decline as a threat to the resilience of carbon stocks in dynamic tropical landscapes. Although global biodiversity conservation efforts often focus on land mammals and birds because of their high levels of decline (*48*), our results underscore the necessity of protecting and restoring them not only for their own sake but for the ecosystem functions that they provide, including climate mitigation. If carefully designed, projects to protect and restore animal biodiversity and landscape connectivity under the Kunming-Montreal Global Biodiversity Framework may not only achieve biodiversity aims but also support the resilience of tropical forests carbon stocks, offering ‘win-wins’ for biodiversity and climate.

## Acknowledgments

We thank the many researchers that collected the field data that were used in the analysis. This publication was made possible by the generous support of the Government of Portugal through the Portuguese Foundation for International Cooperation in Science, Technology and Higher Education and was undertaken in the MIT Portugal Program. This research was supported by a seed award from the MIT Climate and Sustainability Consortium. The Bezos Earth Fund supported S.C.C-P.’s time on this project. This is a contribution of the MIT Terrer Lab.

## Supplementary Materials

### Materials and Methods

Our analysis involved bringing together disparate data types matched to different methods of analysis. The first set of analyses addressed spatial variation in animal-mediated seed dispersal. We then quantified the link between animal biodiversity and forest carbon storage by estimating seed dispersal disruption, modelling its relationship to aboveground carbon accumulation, and using this model to map both carbon accumulation potential and the impact of seed dispersal disruption on carbon accumulation potential. The flowchart in Figure S7 outlines the data and methods used in the latter set of analyses. We describe each aspect in detail below.

#### Prevalence of animal seed dispersal

To assess the global relevance of seed dispersal by animals and evaluate the prediction that tropical regions are most dependent on animal seed dispersal, we estimated the prevalence of animal-mediated dispersal in woody vegetation plots and analyzed its spatial variation. We paired data on seed dispersal mode of plant species with data on plant species’ relative abundance in woody vegetation plots. Using these data, we calculated a weighted, relative metric indicating the prevalence of seed dispersal by animals at each vegetation plot. Then we produced a map of spatial variation in this metric to illustrate variation in the importance of seed dispersal by animals in woody vegetation globally.

To characterize plant species’ dependence on animals for seed dispersal, we first assembled data on dispersal mode from plant trait databases (*49*, *50*) and additional published studies (*17*, *54–66*). Because different sources reported dispersal mode using different categorizations, we recategorized all records of dispersal mode either as abiotic if the seed dispersal process did not involve an animal (e.g., wind, ballistic) or as biotic if it did (e.g., endozoochory, epizoochory). Because multiple records existed for many plant species and sometimes suggested both biotic and abiotic dispersal (e.g., a conifer whose seeds have wings for wind dispersal but also known to be dispersed by seed-caching birds), we treated individual records of biotic dispersal mode as binary (biotic equates to a value of 1 and abiotic equates to a value of 0) and took the average by species. The resulting value aims to characterizes plant species’ dependence on animal seed dispersal, with possible values ranging from zero, meaning the available evidence shows the species is completely abiotically dispersed, and one, indicating full reliance on animals for seed dispersal. This dataset included 346,501 seed dispersal mode records for 24,660 plant species from 4,693 genera and 369 families.

Next, we assessed the prevalence of biotic dispersal in a global database of vegetation plots, sPlotOpen (*67*). Because the focus of the study is on forest regrowth, we examined the subset of plots labelled as containing forest and existing in natural landscapes. This database offers relative abundance measures. We developed a community-weighted measure of biotic dispersal using the biotic dispersal estimates described above and the relative abundance estimates of individual species from the plot data. In each case where dispersal mode was unavailable, we used average values at the finest taxonomic resolution available (e.g., genus, family) to fill missing values. We used the Taxonomic Name Resolution Service (*68*) to harmonize plant species names. Because of an error associated with integration of plots into sPlotOpen from the SALVIAS network (one of many sources of vegetation plot data that are integrated within the larger sPlotOpen dataset), we used the original records of the SALVIAS plots that are also hosted by BIEN, accessing it using the ‘rbien’ package (*50*). In these cases, we calculated an analogous measure of relative abundance in terms of biomass using records of stem diameter, data from BIEN on wood density, and the ‘BIOMASS’ package (*69*) in R, harmonizing species names and using taxonomy to fill in missing wood density data as above.

To evaluate the prediction of higher dependence on animal seed dispersal in tropical forests, we fitted the relationship between latitude and biotic dispersal prevalence with a generalized additive model implemented using the ‘mgcv’ package (*70*). We report average biotic dispersal prevalence among plots in tropical biomes, with biomes defined by Dinerstein *et al.* (*71*). To map biotic dispersal prevalence, we developed a random forest model with the ‘ranger’ (*72*) and ‘tidymodels’ (*73*) packages in R. The biotic dispersal prevalence of each plot was the response variable and the predictor variables were the bio variables from WorldClim version 2.1 at 5 minute resolution. We considered map accuracy by fitting the random forest regression model with three quarters of the dataset (N = 12,793) and withholding a quarter as a test set (N = 4,258), evaluating performance using *R*^2^. During model training, we performed spatially clustered five-fold cross validation implemented using the ‘spatialsample’ package (*74*), testing a grid of hyperparameters. We used a full model using the best-performing hyperparameters to develop the map of biotic dispersal prevalence across both tropical and temperate forest and savanna biomes (Fig. 1). We present model uncertainty as a bootstrapped coefficient of variation by fitting the model over ten resamples of the dataset (Fig. S1).

#### Quantifying seed dispersal disruption

We estimated the severity of disruption of animal-mediated seed dispersal by linking three sets of analysis that describe 1) the effectiveness of each animal species for seed dispersal, 2) the composition of animal species across location effected by habitat fragmentation, and 3) how anthropogenic alteration of animal movement alters seed dispersal. We develop these estimates for birds and mammals, which are the groups with the major role in animal-mediated primary seed dispersal across most ecosystems and those with the highest data availability. Other animal groups of seed dispersers can play important roles, such as the important role of reptiles in island ecosystems, and we assume that the broad patterns of seed dispersal disruption that we identify largely apply across these other groups as well. Below we describe the data used to characterize these three processes and finally how we bring them together to estimate a metric of seed dispersal disruption (Fig. S7).

##### Seed dispersal effectiveness

We incorporate species-specific estimates of seed dispersal function by birds and mammals developed by Fricke *et al.* (*28*). Briefly, their analysis involved a large synthesis of published studies of the seed dispersal process and trait-based machine learning models to predict seed dispersal function for each species. The training data they assembled from the literature included seed dispersal records of 406 local ecological networks, data on seed retention times and movement across 302 animal species, and results of 2215 studies of the impact of gut passage on seed germination. Traits of birds and mammals include traits describing their morphology, life history, and diet (*75*). Their trait-based approach to model stages of the seed dispersal process for individual species showed high cross-validated performance on a withheld tenth of the data; for example, in predicting which animal species disperse seeds (AUC = 0.90), seed retention times (*R*^2^ = 0.96), dimensions of animal movement (*R*^2^ = 0.80, *R*^2^ = 0.73), and germination rates following gut passage (*R*^2^ = 0.36) (*28*). In the analysis of Fricke *et al.* (*28*), the outputs of these models were combined (overviewed in panels 1 and 2 in Fig. S7) to generate a species-specific measure of seed dispersal function above a given distance threshold from a mother plant of a hypothetical representative plant species. The metric is unitless but is proportional to the number of seeds dispersed past the distance threshold that germinate per unit time. This approach thus emphasizes the quantity of effective seed dispersal, which is important for overcoming the seed limitations that often hampers regrowth (*13*). By following the 1 km threshold of long-distance dispersal defined in ref. (*28*), this approach characterizes relatively long-distance seed movements that are important for plant recovery following disturbance (*11*, *29*).

##### Animal composition

To incorporate information on which species provide seed dispersal function at a location, we first considered data on species ranges provided by the IUCN/BirdLife International. We used range data for extant species in their current ranges. While range information outlines where species can occur, whether species occur at individual sites is also determined by local processes, and we focus on how habitat loss and fragmentation impact animal populations locally. To incorporate this effect, we used data from a meta-analysis of records of local occurrence in fragmented forest landscapes (*76*). The data provide information on the probability that species that exist in the region occur at specific locations, which vary in the levels of fragmentation. Although the dataset focuses on birds, quantitatively similar patterns are known from studies of mammals (*77*) for which similar global syntheses are unavailable. To characterize habitat fragmentation, we used a simple measure of proportional tree cover per area in a 500m radius circle around the observation location from the local occurrence dataset. We calculated this using Earth Engine implemented in the ‘rgee’ package (*78*) in R with tree cover data (*51*) from the year 2000. Although the occurrence data spans decades, this is roughly the average year of field recording in this dataset. The simple metric of forest habitat fragmentation enables us to characterize fragmentation for the sites of field records and to apply pixel-wise calculations during the map development described below. In a generalized linear mixed effects model, the binomial response variable reflected species presence or absence and the predictor variable was the measure of forest habitat fragmentation. We included random intercepts for biome, species name, and study ID. We used the fitted relationship (Fig. S3A) along with species composition for a location based on range data to calculate a weighted occurrence value for each species describing their density on the landscape (*79*).

##### Seed disperser movement

Seed dispersal function also depends on landscape factors that alter animal movement; the movement of individual animals is affected by anthropogenic impacts on landscapes (*19*), and this in turn influences seed dispersal distances (*20*). To incorporate these effects into our estimates of seed dispersal disruption, we repeated the approach applied by Tucker *et al.* (*19*). This approach involved taking animal movement records and calculating displacement (the distance between an individual’s location at one time and another) over time from an arbitrary starting location, and then relating displacement to a holistic metric of anthropogenic environmental impact, the human footprint index (HFP) (*53*). We incorporated data provided on MoveBank under CC0 licenses (Table S2) for terrestrial mammals and birds. Using the movement records, we randomly chose timestamped records and used subsequent timestamped location records over a set time period to represent a hypothetical seed dispersal event. The time periods over which we developed these movement tracks were chosen to encompass typical seed retention times given species’ traits based on modelling by ref. (*28*). We allowed up to five non-overlapping periods of these hypothetical seed dispersal events per animal individual, leading to 7769 periods total. For movement tracks with frequent location records, we thinned this frequency to improve computational efficiency, although this did not quantitatively affect results. Following Tucker *et al.* (*19*), we extracted HFP for each location record and also included remotely sensed NDVI as a covariate. Using the MOD13Q1.061 Terra Vegetation Indices dataset from the Earth Engine data catalogue accessed via the ‘rgee’ package, we used average monthly values for the corresponding month over the period 2001 to 2021, and otherwise associated individual movement records with the nearest corresponding calendar month with available data. In a general linear mixed effects model, log-transformed displacement values were the response variable, and the predictor variables were log-transformed time and average values for each of HFP and NDVI. To account for sources of non-independence, we allowed random slopes and intercepts by species and random intercepts by time period. The fitted model (Fig. S3B) was used to estimate the proportional reduction in seed dispersal distance caused by a given level of HFP.

##### Calculating seed dispersal disruption

To assess the severity of seed dispersal disruption, we compared current levels of seed dispersal to levels of seed dispersal in a scenario of seed dispersal not disrupted by anthropogenic landscape impacts. We first calculated current seed dispersal function by modifying the seed dispersal estimates as calculated by Fricke *et al.* (*28*) by incorporating the anthropogenic impacts on species presence and movement outlined above. In the scenario without seed dispersal disruption, we similarly calculated seed dispersal function as a present counterfactual lacking habitat fragmentation and with HFP equal to zero. We calculated seed dispersal disruption as the proportional decline between the current values and the values without the drivers of seed dispersal disruption. This gives seed dispersal disruption as a unitless value bounded between zero (no seed dispersal disruption) and one (complete seed dispersal disruption). We used seed dispersal disruption values associated with the tropical regrowth locations described below and in analyses that involved mapping seed dispersal disruption across the tropics, which we calculate at ∼1 km resolution. When expressed in a Behrman equal area projection for visualization and analysis, pixels represent *c.* 0.71 km^2^. To estimate seed dispersal disruption in 2020, we repeated the above steps to calculate seed dispersal disruption, but incorporating tree cover changes between 2000 and 2020 (*52*) to update estimates of habitat fragmentation. We use both the circa 2000 and 2020 estimates of seed dispersal disruption in our analyses below.

#### Modelling aboveground carbon accumulation

##### Forest carbon data and covariates

To assess how forest regrowth over time is related to seed dispersal disruption and variables related to forest growth, we used field records of the natural regrowth of aboveground biomass or carbon per hectare over stand age assembled by Cook-Patton *et al.* (*26*). We analyzed the portion of these data from the tropics (*71*) and converted biomass values to carbon units by multiplying by 0.47 following ref. (*80*). To help disentangle the relevance of seed dispersal disruption on observed forest regrowth differences, we also include in the analysis data from a corresponding dataset of field records of carbon accumulation in monoculture plantations assembled by Bukoski *et al.* (*32*). We include the tropical aboveground carbon per hectare records available in this dataset. Monoculture plantation sites show similar ranges of environmental variables as observed in the natural regrowth sites (Fig. S8) but were more prevalent at higher latitudes and somewhat cooler and drier locations. We analyzed resurvey data (that is, after stand age of zero), including 2014 natural regrowth records and 1351 records from monoculture plantations. In the analysis, we used seed dispersal disruption values estimated circa 2000, which is roughly the average year across records where date information is available.

We included additional variables that can influences forest growth or that can pose limitations on regrowth in anthropogenically modified landscapes: temperature, precipitation, drought, fire, grazing livestock, and potential net primary productivity. We used WorldClim annual mean temperature and total annual precipitation values. To directly model constraints on regrowth caused by heat- and desiccation-related edge effects, we used the additive inverse of the Vegetation Health Index (*81*) as a descriptor of drought conditions that can occur within fragmented landscapes. We used averaged values of the monthly VHI values provided by UNFAO at 1 km resolution over the period 2001-2011. We masked to areas that maintained at least 10% forest cover (*51*) over this period to capture drought condition of (regrowing) forest areas in the local area rather than of other land cover types. We calculated average values within a 2500 m radius, or within 5000 m if data were otherwise missing. Fire also limits forest regrowth, and we used the MODIS Fire-cci Burned Area version 5.1 product at 250 m resolution to characterize the prevalence of fire on the landscape, calculated number of fire days for each pixel, and masked and averaged within a buffer as above. Although we calculated these metrics over the period 2001-2011, the values were highly correlated with other time periods (e.g., *R*^2^ > 0.99 when correlated with values for the period 2011-2020). We incorporated information on the density of grazing animals, which may also limit regrowth in anthropogenically modified forest landscapes. Using data from the Gridded Livestock of the World (GLW 3) product (*82*) estimated *c*. 2010 at 10 km resolution, we summed the abundance of cattle, sheep, and goats and extracted values at the site coordinates. To capture the net residual effect of additional factors that constrain or favor forest growth in a given landscape, we included potential net primary productivity as a predictor variable. Because anthropogenic land use often reduces net primary productivity (NPP) (*83*), directly observed NPP values for a given location may not accurately reflect the NPP that would occur if forests were regrowing at that location. We therefore aimed to estimate potential NPP using observations of NPP from nearby areas of natural vegetation. To do so, we started with the MOD17A3HGF.061 Terra Net Primary Production dataset accessed via Earth Engine and the ‘rgee’ package in R. We calculated potential NPP by first averaging NPP values over the period 2001-2011 and then calculating the maximum value over a 10 km radius. This has the effect of smoothing out local reductions in NPP caused by anthropogenic land use, offering a proxy for the maximum potential NPP that would occur in a pixel if it were covered by natural forest vegetation. The potential NPP values thus enable the modelling of overall site productivity influenced by landscape-scale drivers not captured directly by other predictor variables. This could include factors such as soil degradation or forest resource extraction for which direct pantropical estimates are not available.

##### Forest carbon accumulation model

In a Bayesian model implemented in JAGS using the ‘rjags’ package (*84*) in R, we fitted a saturating model of aboveground carbon (*μ*) over time in years (*t*) using the Monod equation.

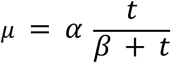

The parameters describe peak carbon accumulation (*α*) and the half-saturating time (*β*). We allowed *α* to vary with key factors related to maximum forest carbon: temperature, precipitation, and potential NPP. We allowed *β* to vary with these variables as well as potential constraints on regrowth: seed dispersal disruption, fire, drought, and grazers as described above. For both *α* and *β*, we fitted a site-specific ‘random effect’ to account for non-independence among the multiple observations per site and for other spatial variation not captured by the predictor variables. To examine our prediction that in monoculture plantations seed dispersal disruption would have no negative impact on forest growth, we included an interaction term so that records from monoculture plantation and natural regrowth plots could exhibit different relationships to seed dispersal disruption. For monoculture plantation sites, we additionally included a coefficient to describe the effect of additional interventions of fertilizing, weeding, irrigation when these interventions were reported (*32*). In all cases, we used weakly informative priors and sum-to-zero constraints for the site-level effects. We ran three chains for 500,000 iterations, sampling with a 500-iteration thinning interval after 10,000 iterations each of adaptation and burn-in. We developed posterior predictions for forest growth over time for varying levels of seed dispersal disruption for both monoculture plantation and natural regrowth. For results presented as accumulation rates, we calculated an annualized accumulation rate over the first 30 years of regrowth. We also calculated a standardize effect size of the predictor variables; the units of the effect size represent the difference in carbon accumulation rate caused by a 1 standard deviation difference in the predictor variable, holding other variables at their average. We report 95% credible intervals for statistical inference. We also implemented a second version of the analysis that was otherwise identical but did not estimate the effects of seed dispersal disruption to compare model estimates with and without considering seed dispersal disruption. Full details of model implementation are presented with the data and code package.

#### Mapping estimates and uncertainty

We developed maps of carbon accumulation potential presented as an annualized rate based on mean posterior values from the fitted model of forest growth and maps of each predictor variable. For the current estimate, we used the map of seed dispersal disruption estimated circa 2020. We assessed how changes to seed dispersal disruption since 2000 have affected regrowth potential by taking the difference between current values and estimates based on seed dispersal disruption circa 2000. Finally, to estimate how much carbon accumulation potential has been lost due to seed dispersal disruption, we performed otherwise similar calculations but set seed dispersal disruption to zero and took the difference with the current map of regrowth potential.

To characterize uncertainty, we report the magnitude of the 95% credible interval of accumulation rates. To do so, we took 100 samples of the posterior distribution for each coefficient, mapped carbon accumulation rate at each pixel as above, and reported the difference between the 2.5 and 97.5 percentiles. We reported an equivalent uncertainty measure for the lost carbon accumulation potential attributable to seed dispersal disruption. To examine the influence of seed dispersal disruption on natural regrowth within sites where natural regrowth may be implemented for ecosystem restoration and climate mitigation, we considered the map of restoration areas developed by Griscom *et al.* (*36*). We present lost potential of accumulation rates in these areas by estimating carbon accumulation if seed dispersal were not disrupted versus accumulation rates under current levels of seed dispersal disruption. To characterize uncertainty in these estimates, we present a magnitude of uncertainty as described above, summing pixel-wise the low estimates (2.5 percentile of accumulation rate at each pixel) and high estimates (97.5 percentile) and then taking the difference. Note that our carbon accumulation estimates do not include belowground carbon accumulation, nor soil carbon accumulation, associated with natural regrowth.

**Fig. S1.**
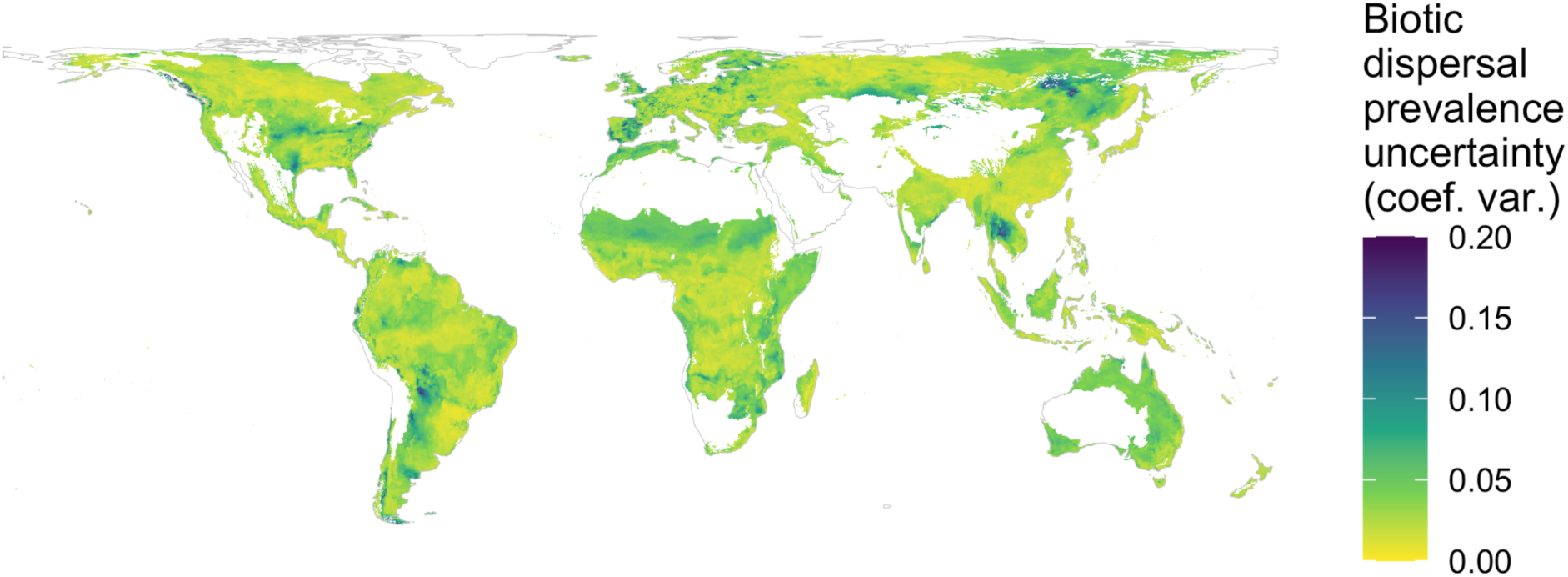
Model uncertainty for the prevalence of biotic seed dispersal in woody vegetation across forest and savanna biomes expressed in terms of the coefficient of variation across models refitted using bootstrapping.

**Fig. S2.**
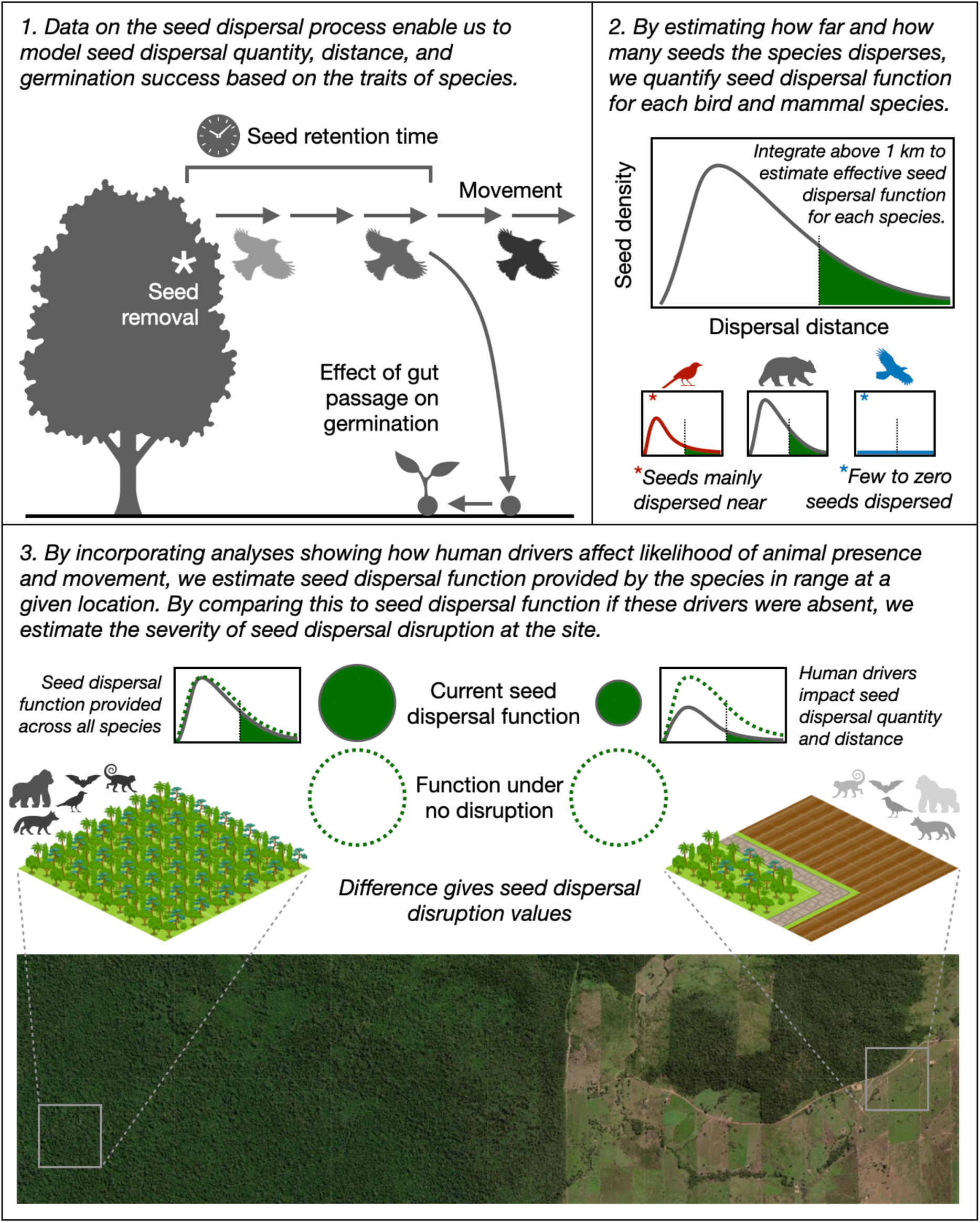
Overview of approach to estimate seed dispersal function and the severity of seed dispersal disruption. See Fig. S7 for overview of datasets used to determine seed dispersal disruption.

**Fig. S3.**
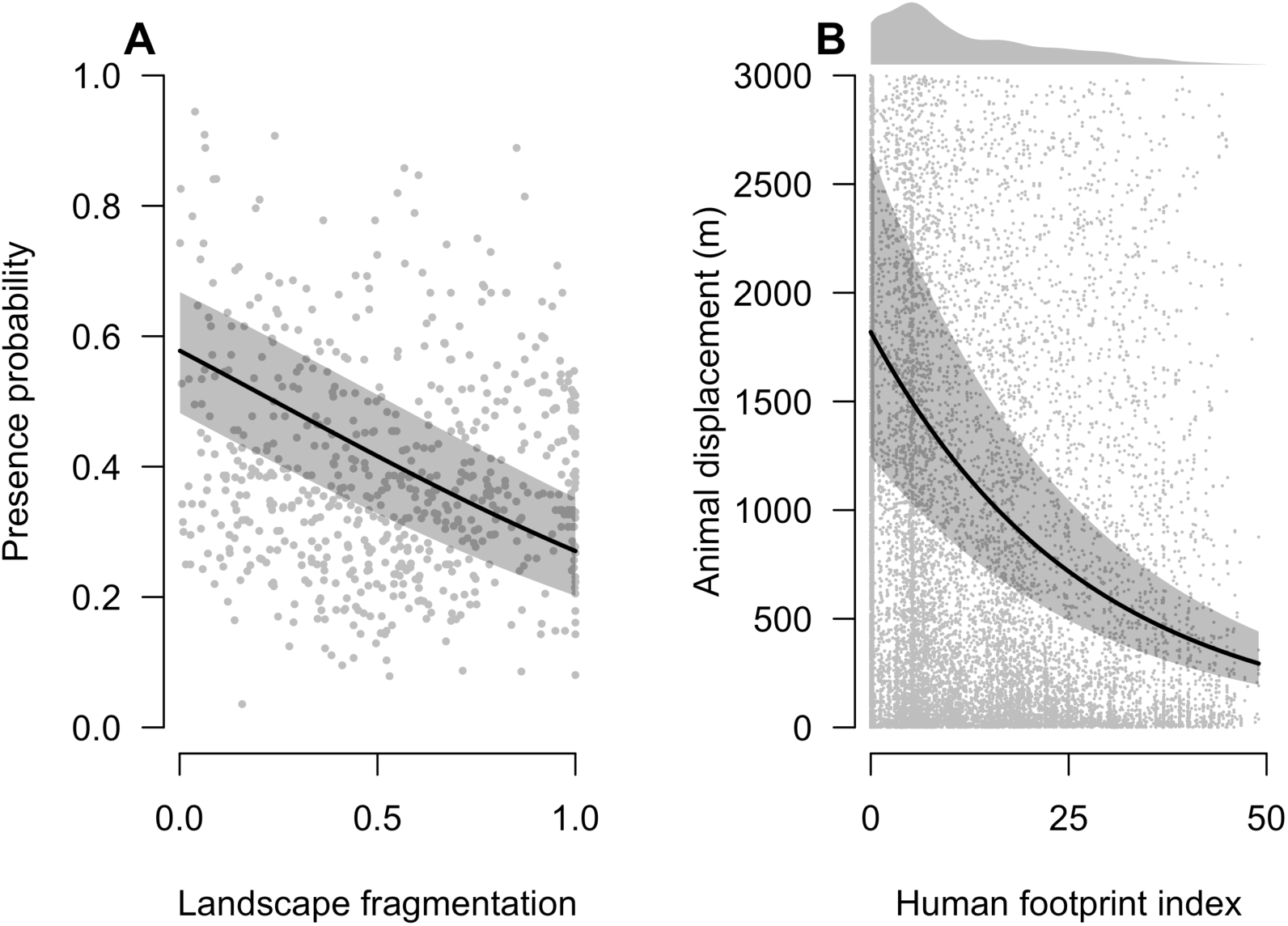
Model results for the effect of land use on the presence and movement of animals. (**A**) Forest species presence was significantly related to landscape fragmentation measured as proportional tree cover (generalized linear model standardized coefficient: −1.31 ± 0.06 S.E.; *P* < 0.0001; n = 61,716). Points show average presence at each study site for data visualization. (**B**) Animal displacement distances were significantly related to the human footprint index (generalized linear model standardized coefficient: −1.86 ± 0.12 S.E.; *P* < 0.0001; n = 488,583; shown for an average time since start of seed retention period). Points are a random sample of three values per tracking period for data visualization, with distribution of displacement values above 3000 m represented by marginal density plot.

**Fig. S4.**
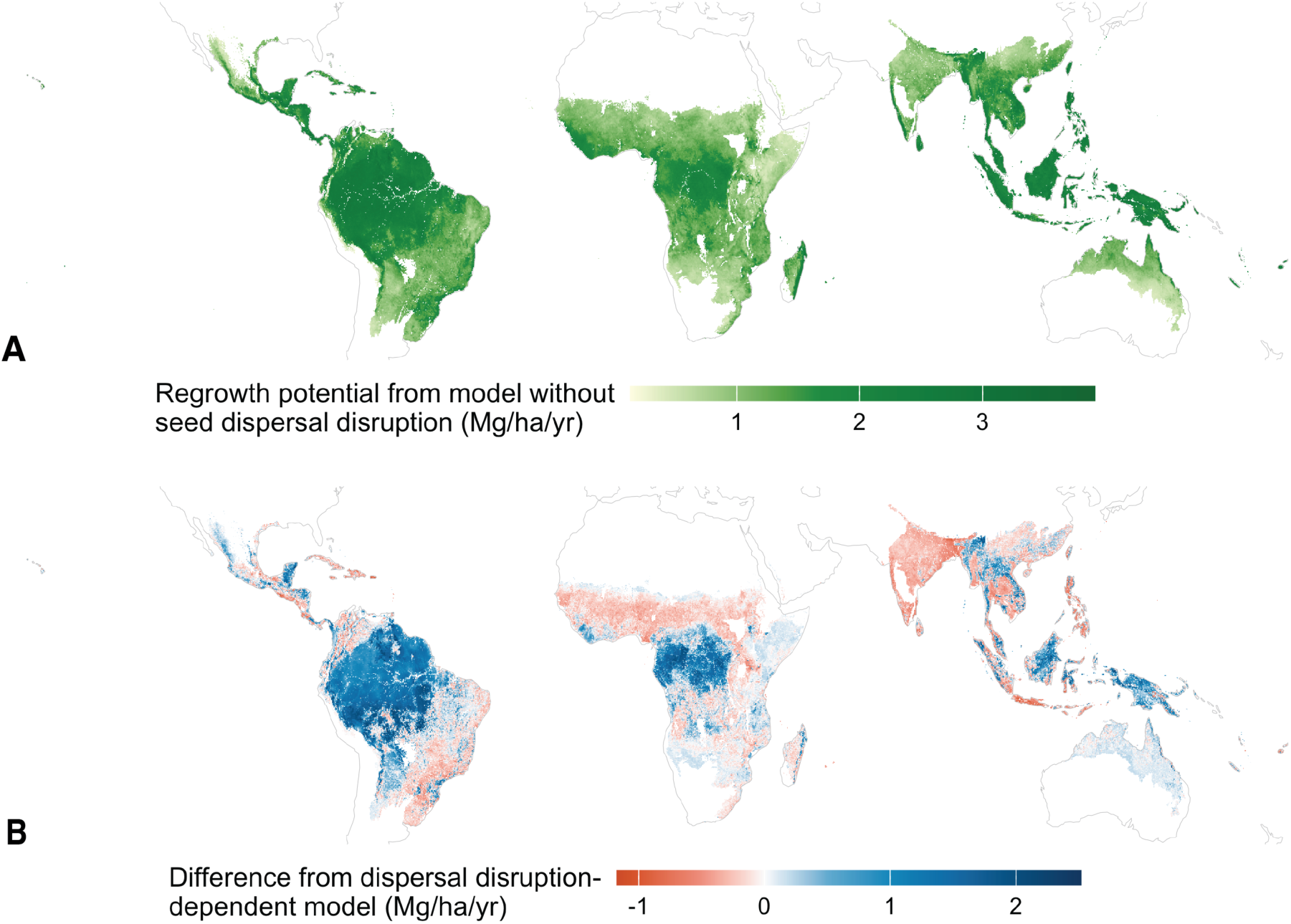
Comparisons to a model fitted without considering seed dispersal disruption. (**A**) Regrowth potential estimated from an otherwise identical model that excluded seed dispersal disruption. (**B**) difference between model fitted with and without considering seed dispersal disruption. Negative values show areas where the model fitted using seed dispersal disruption estimates lower regrowth values than in the model driven only by other environmental variables.

**Fig. S5.**
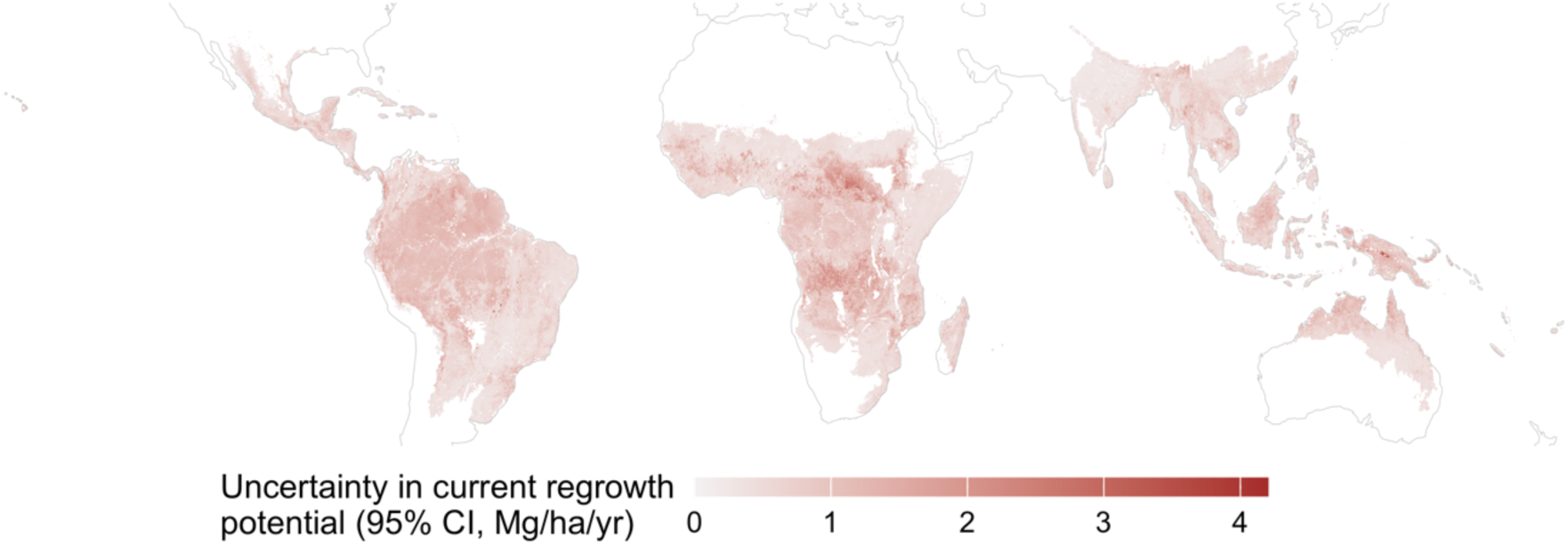
Uncertainty in current regrowth potential, corresponding to median estimates presented in Fig. 4A, measured as the magnitude of the 95% credible interval.

**Fig. S6.**
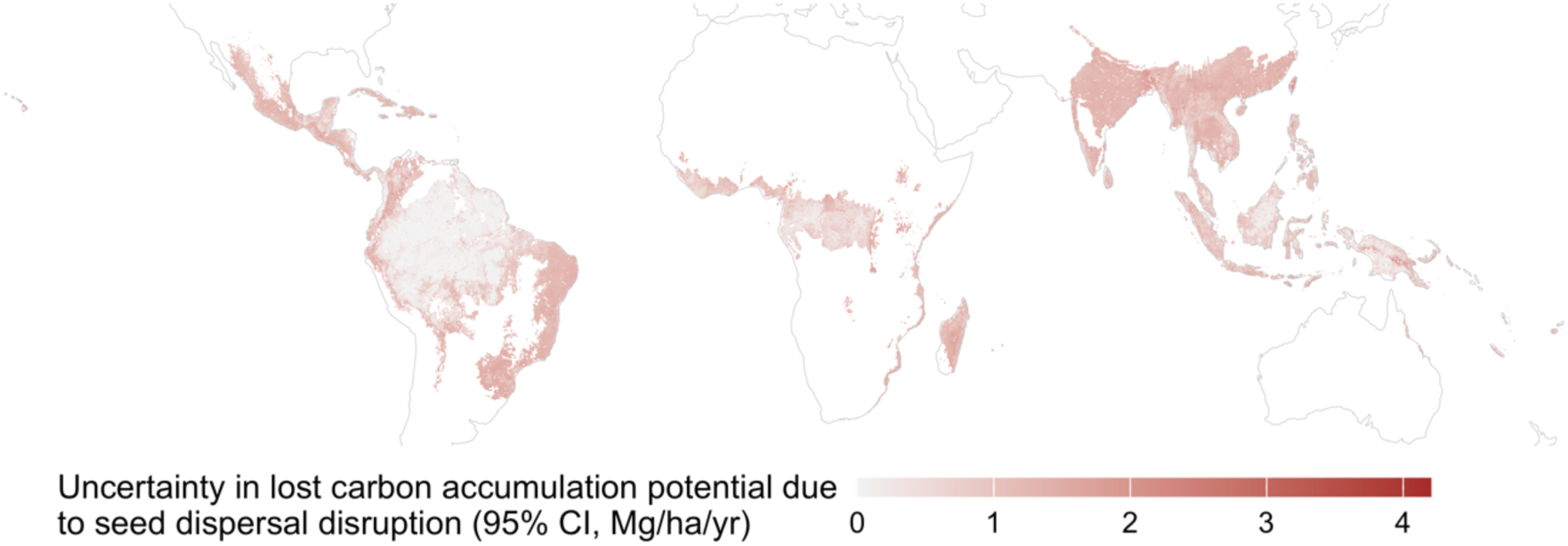
Uncertainty in the lost carbon accumulation potential due to seed dispersal disruption, corresponding to median estimates presented in Fig.

**Fig. S7.**
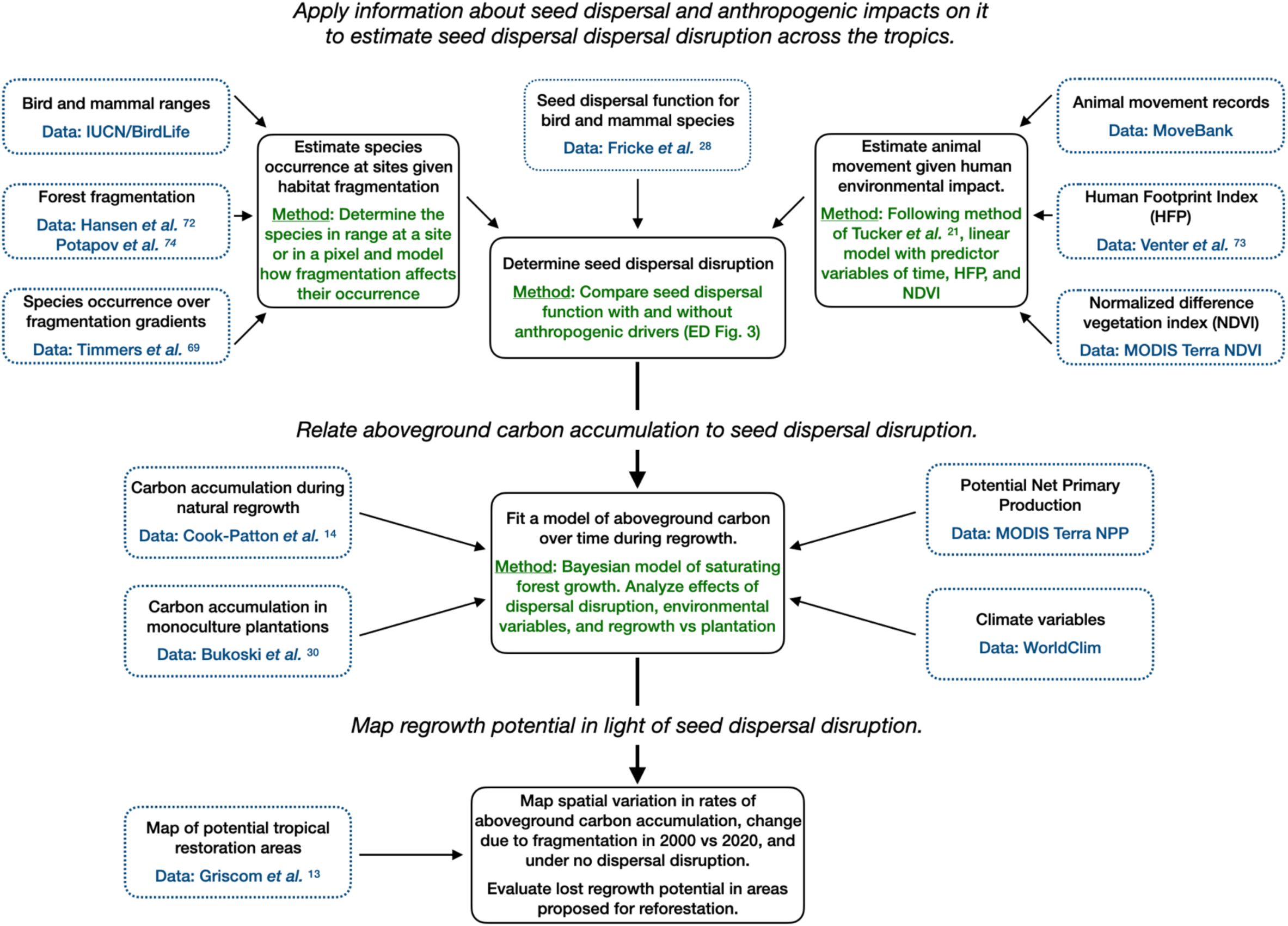
Overview of data and methods used to estimate seed dispersal disruption, its influence on forest regrowth, and the consequences for estimates of climate mitigation potential.

**Fig. S8.**
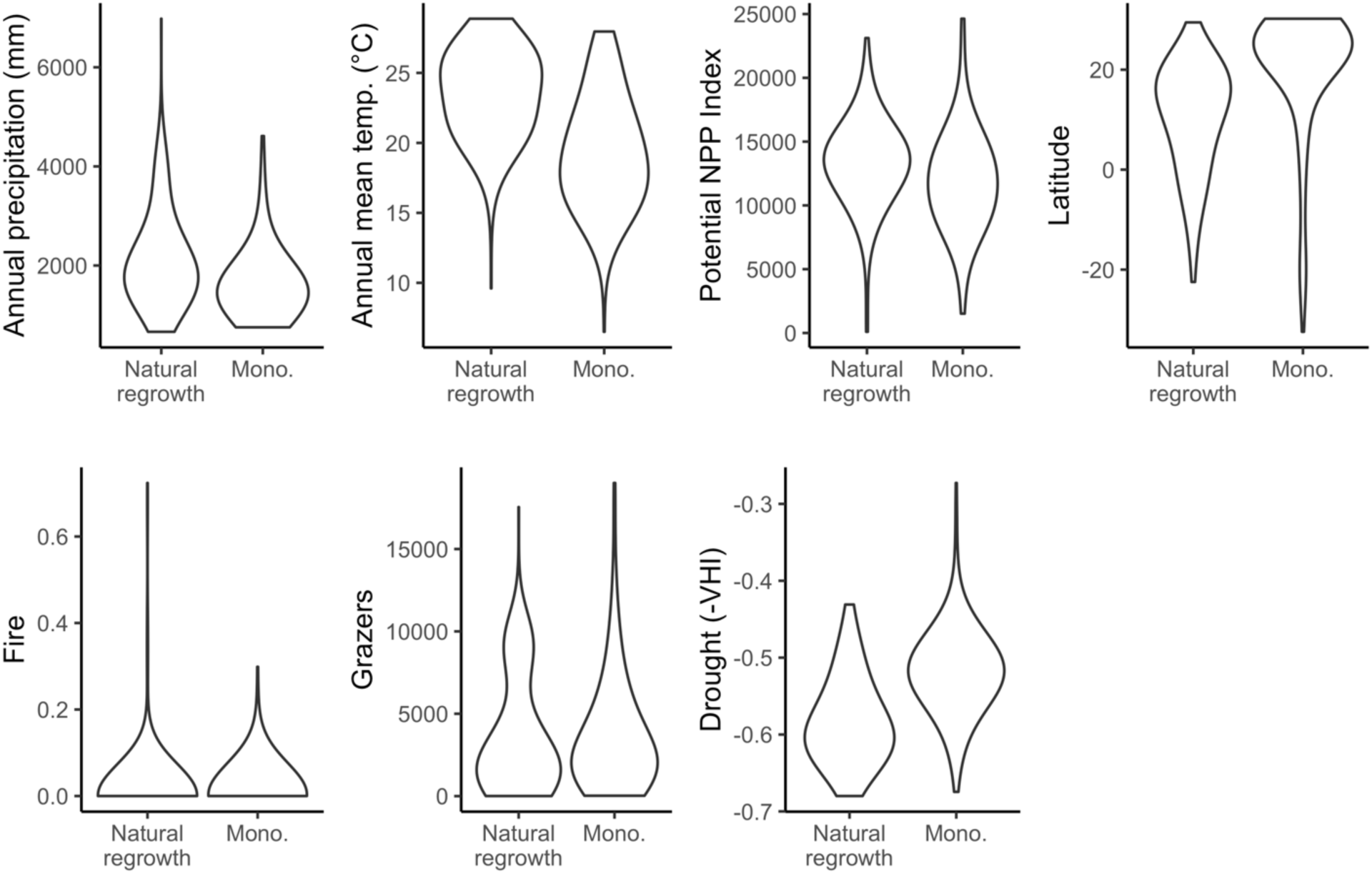
Comparison of the range of environmental variables for natural regrowth and monoculture sites.

**Table S1.** Posterior values of standardized effect sizes, showing the effect on aboveground carbon accumulation annualized over the first thirty years of growth. Columns give the median estimate or quantiles that make up the 95% and 99% credible intervals. Values represents the effect of a one standard deviation increase in the predictor variable or, for the binary plantation intervention term, the use of plantation interventions.

| Coefficient | Median | 0.005% | 0.25% | 0.975% | 0.995% |
| --- | --- | --- | --- | --- | --- |
| Potential net primary productivity | 0.131 | 0.0331 | 0.0547 | 0.21 | 0.249 |
| Annual mean temperature | 0.203 | 0.0452 | 0.0764 | 0.337 | 0.381 |
| Total annual precipitation | 0.288 | 0.138 | 0.169 | 0.432 | 0.471 |
| Fire prevalence | -0.104 | -0.217 | -0.19 | -0.00889 | 0.0204 |
| Grazing livestock abundance | -0.0424 | -0.148 | -0.123 | 0.0444 | 0.0789 |
| Drought | -0.185 | -0.3 | -0.273 | -0.105 | -0.0841 |
| Dispersal disruption (nat. regrowth) | -0.38 | -0.49 | -0.462 | -0.303 | -0.285 |
| Dispersal disruption (monoculture) | 0.153 | 0.0348 | 0.0626 | 0.272 | 0.317 |
| Plantation interventions | 0.925 | 0.493 | 0.575 | 1.38 | 1.56 |

**Table S2.**
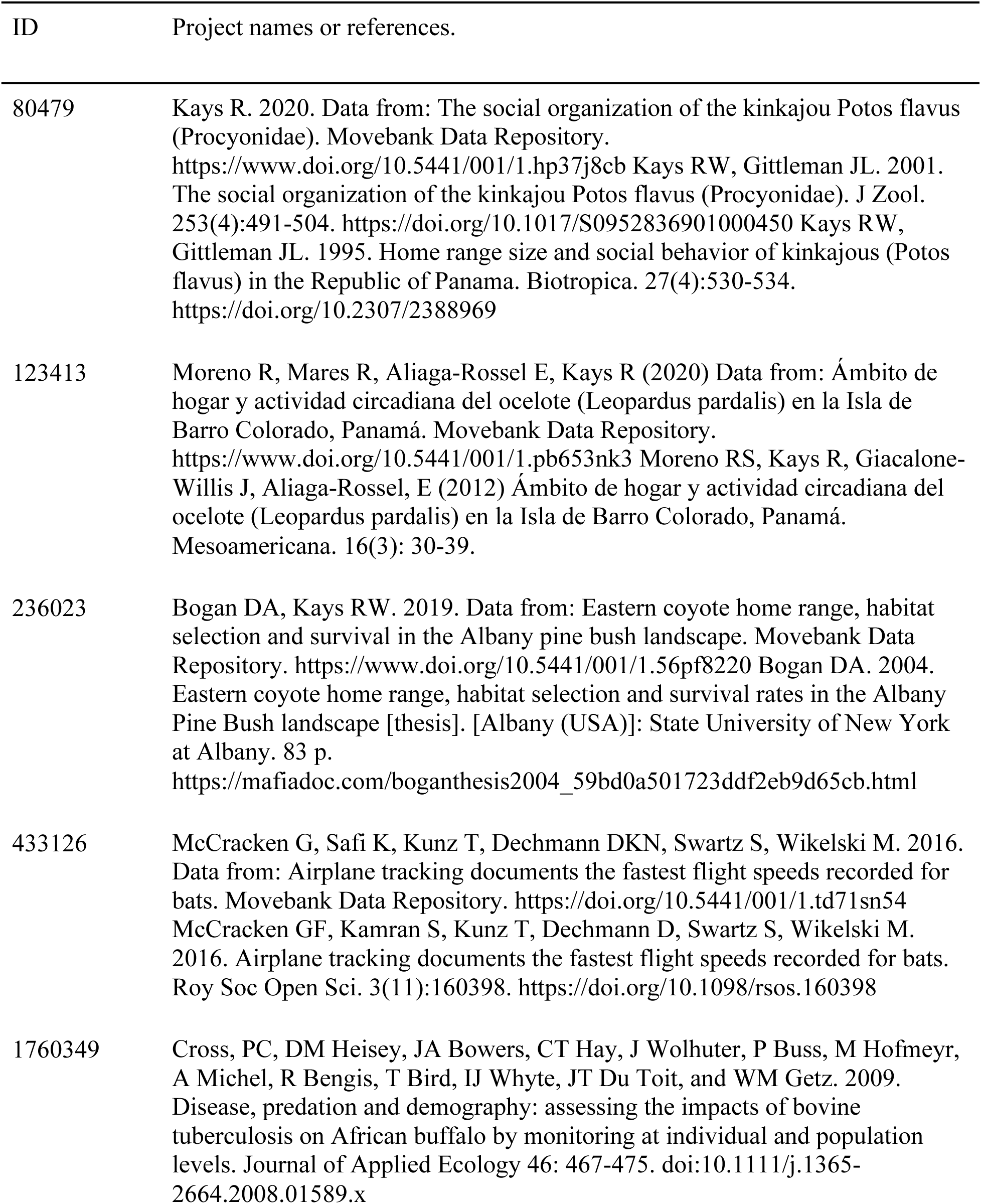
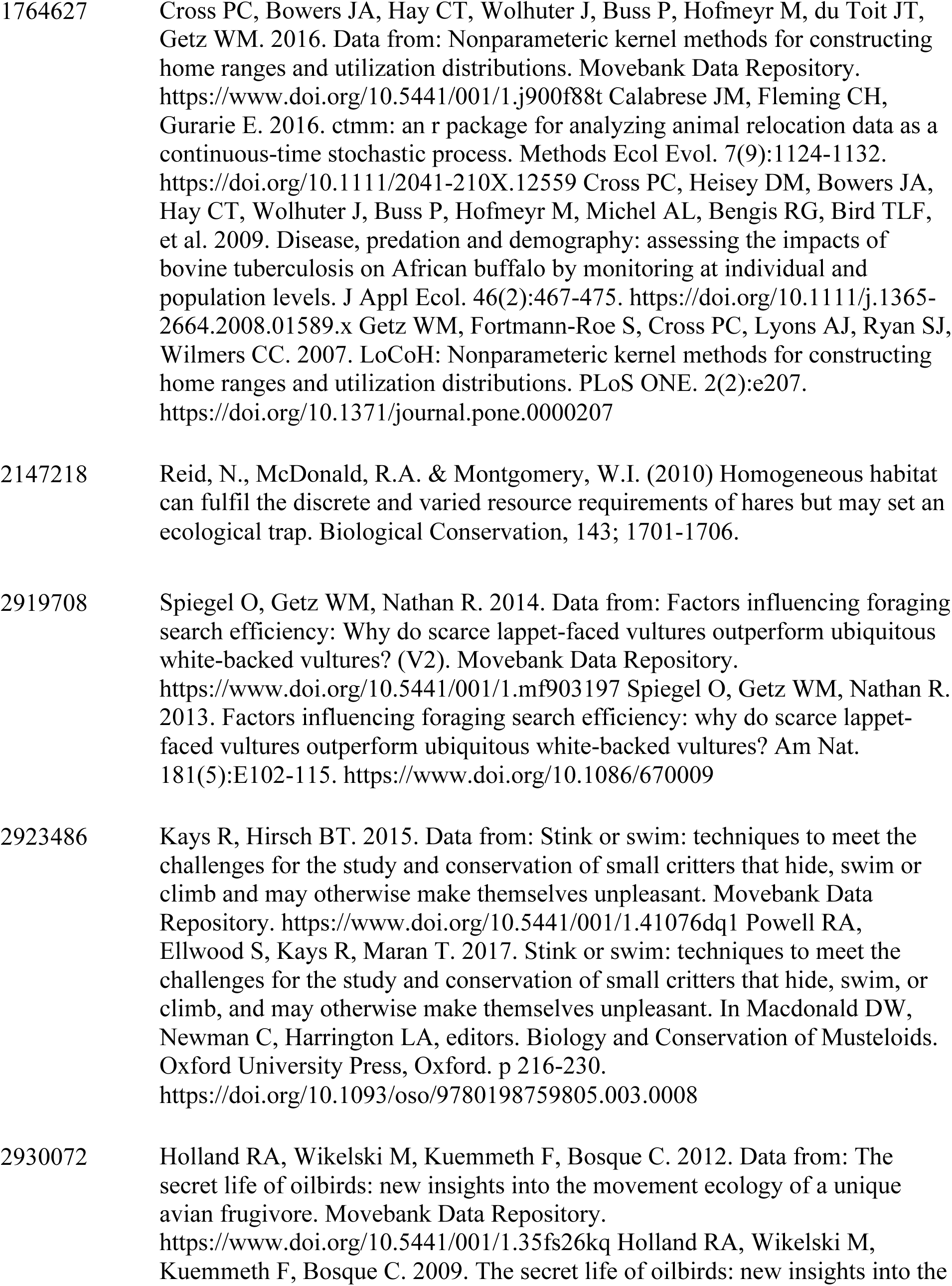
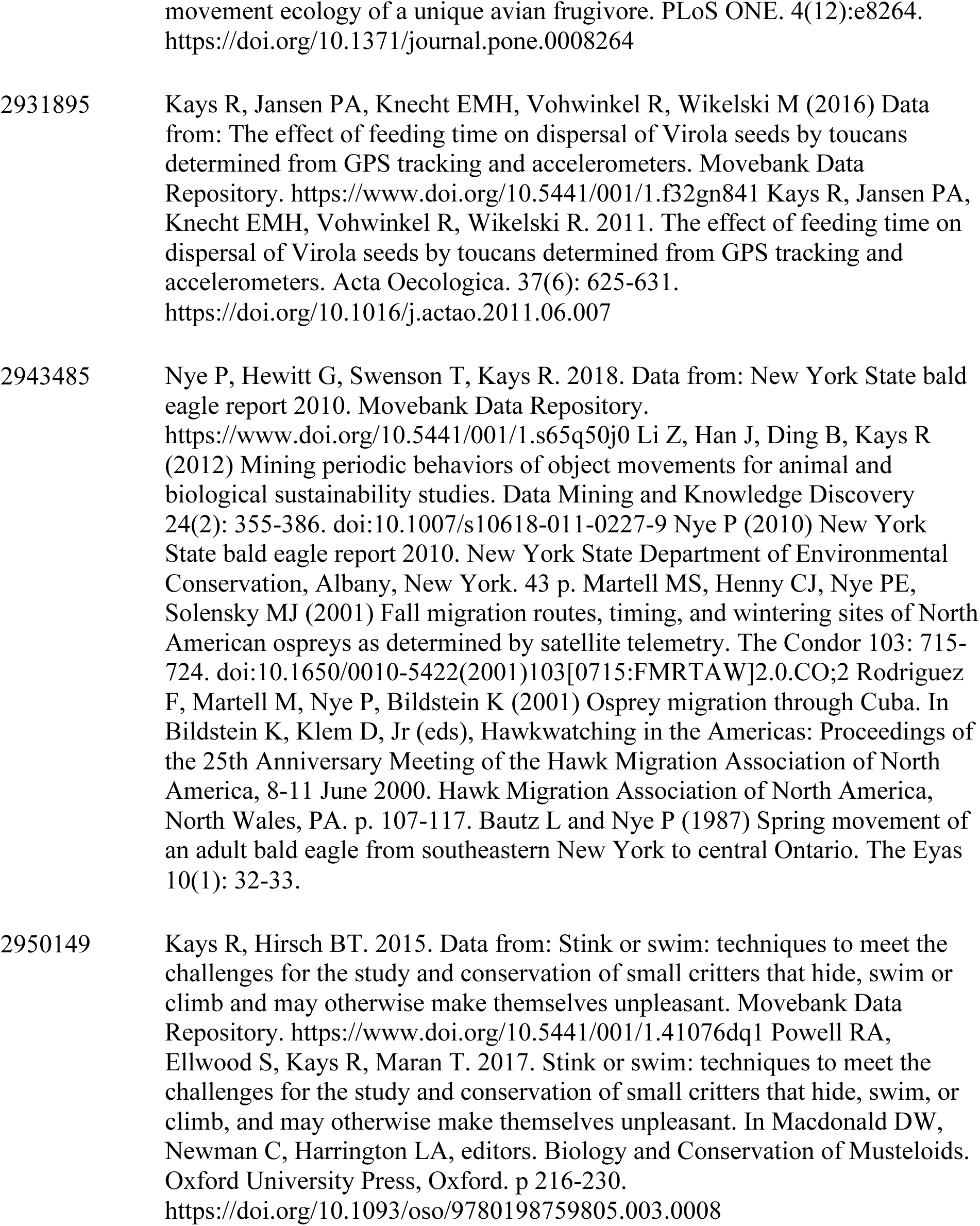
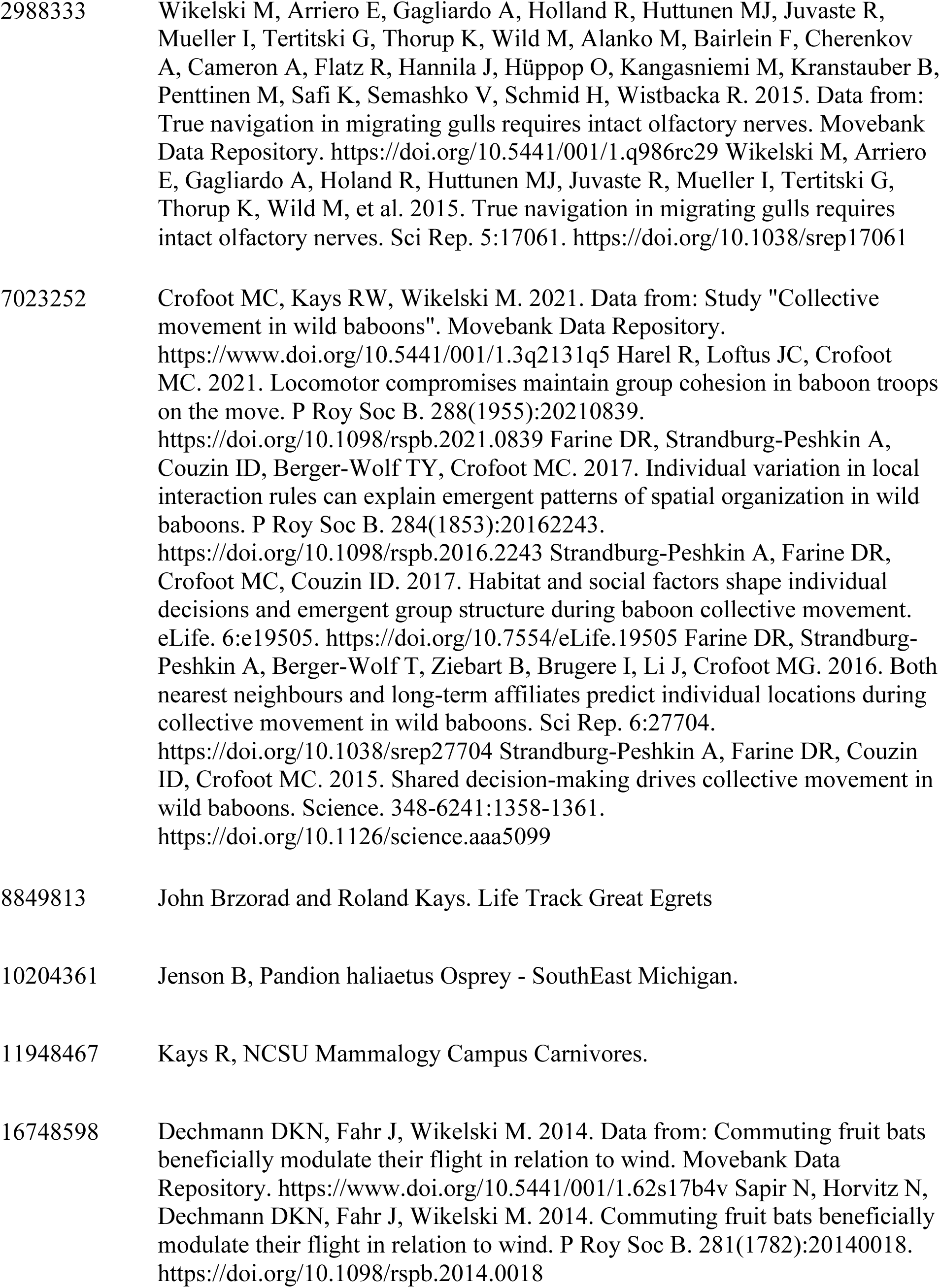
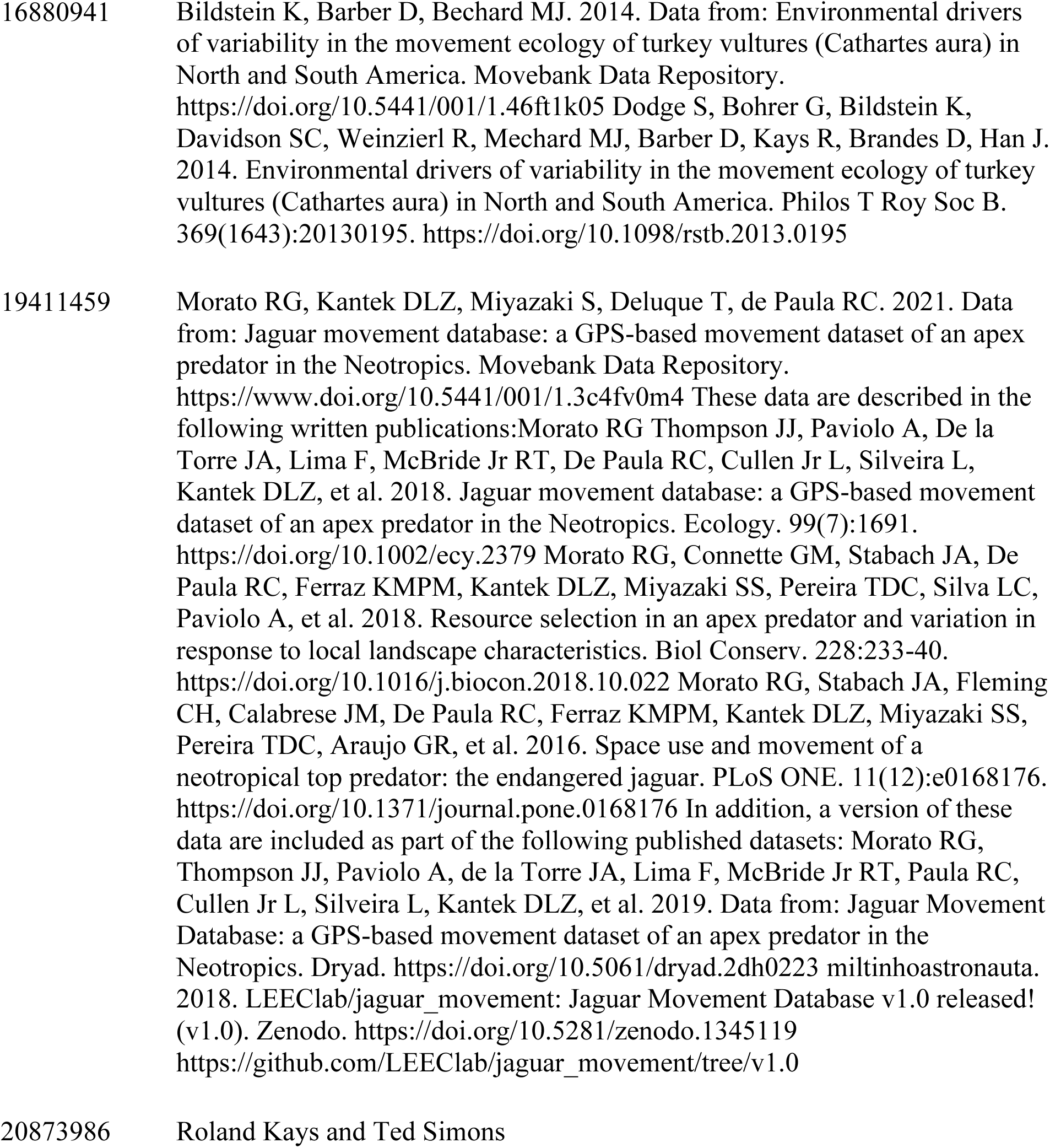
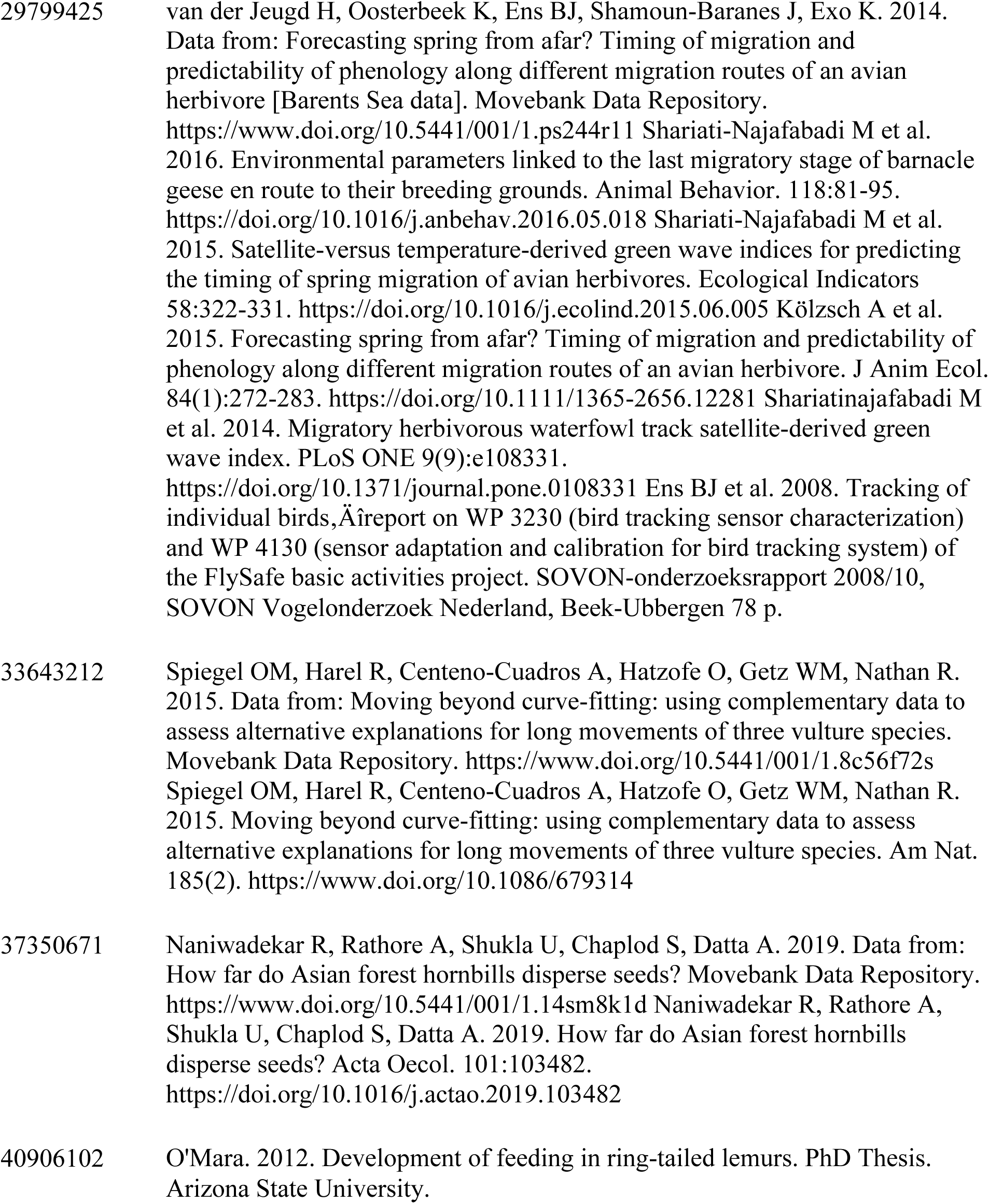
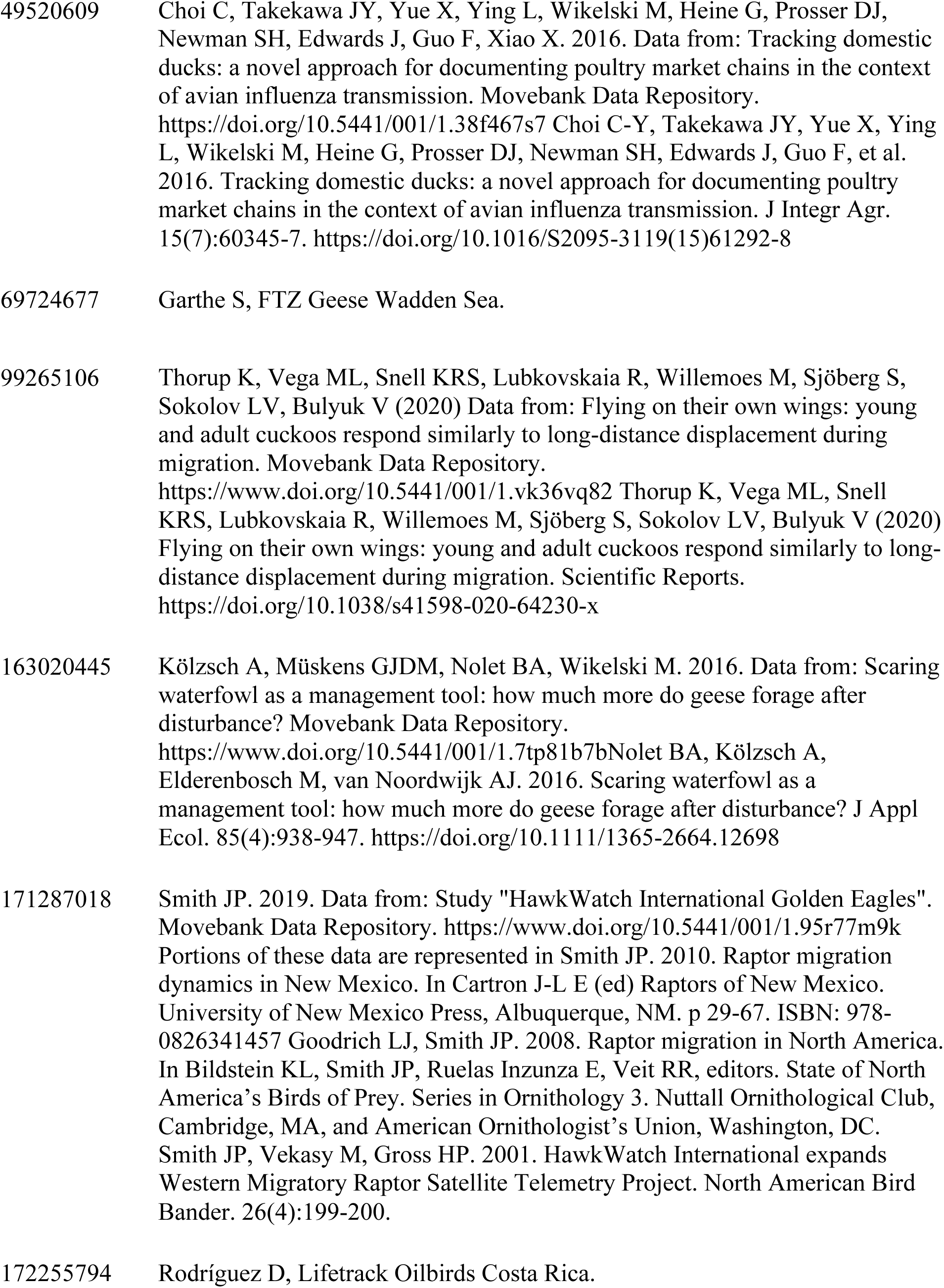
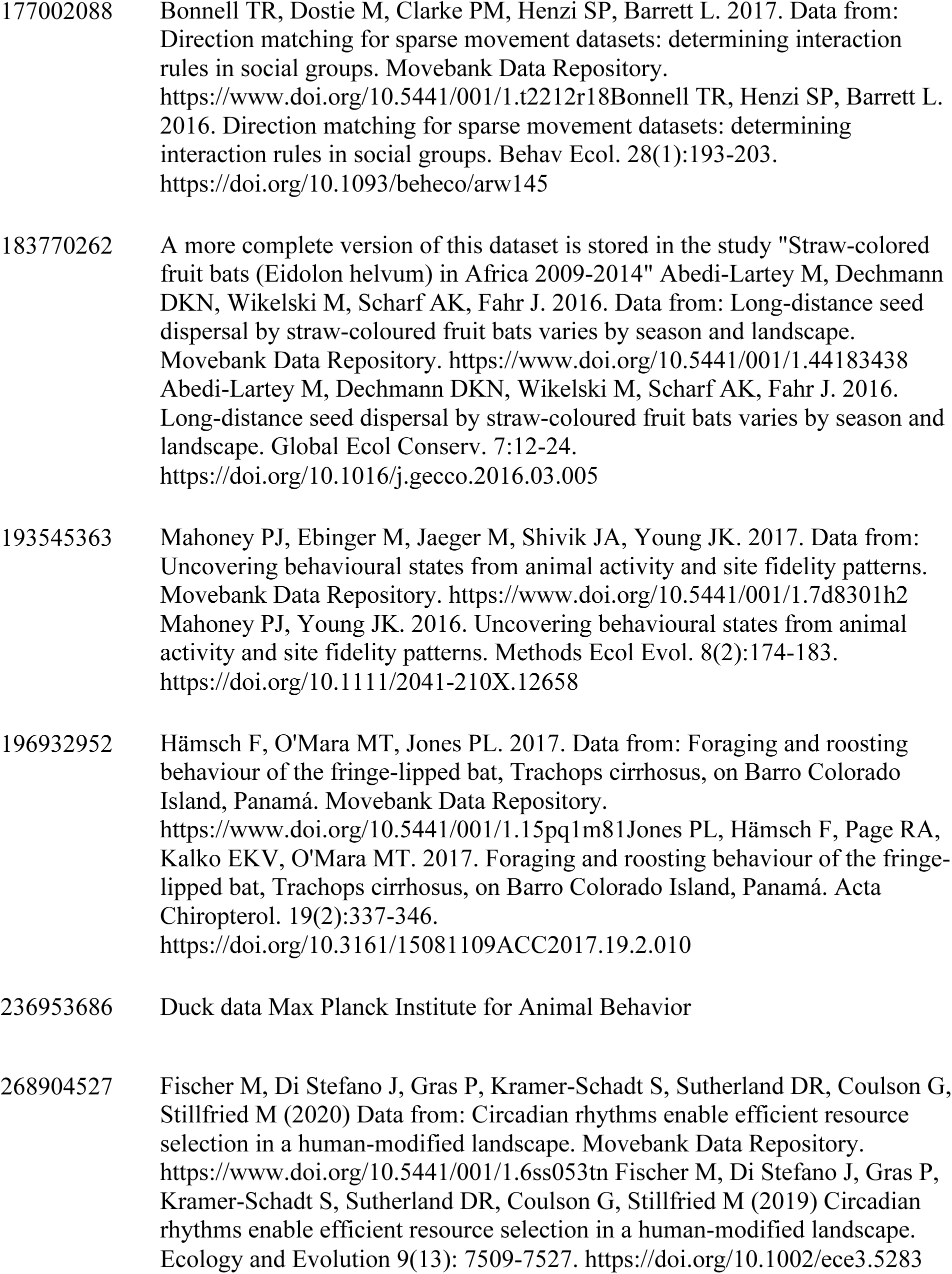
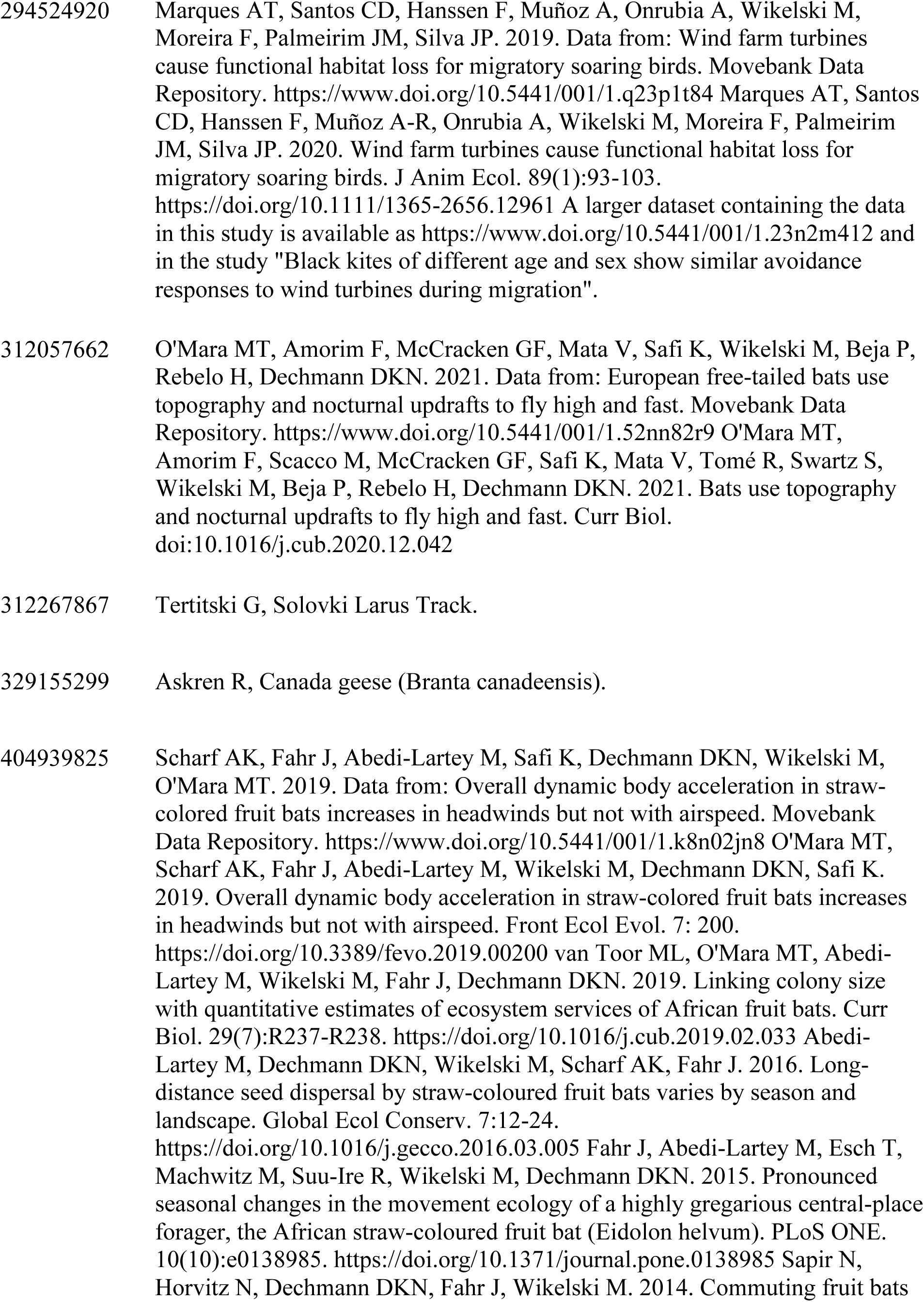
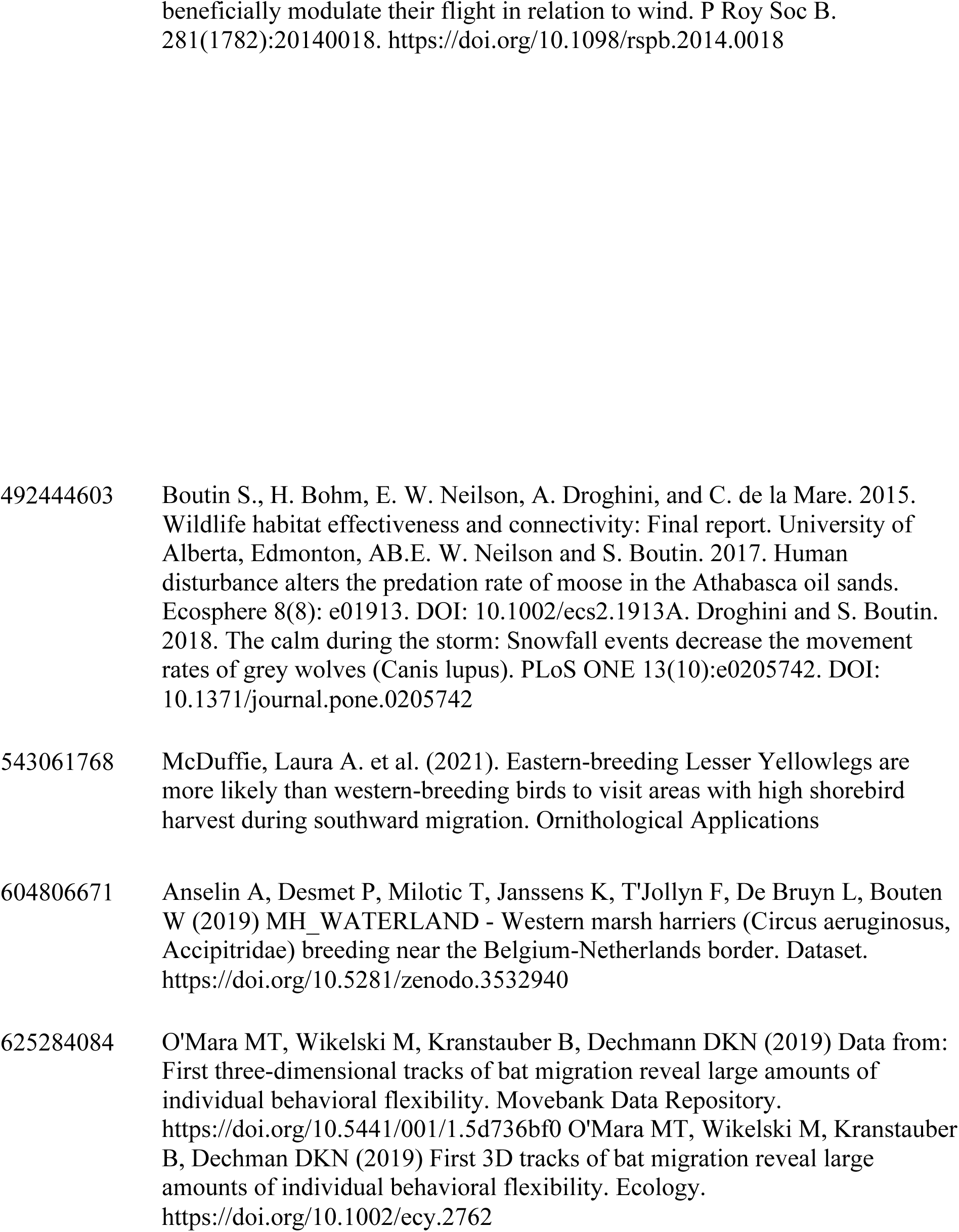
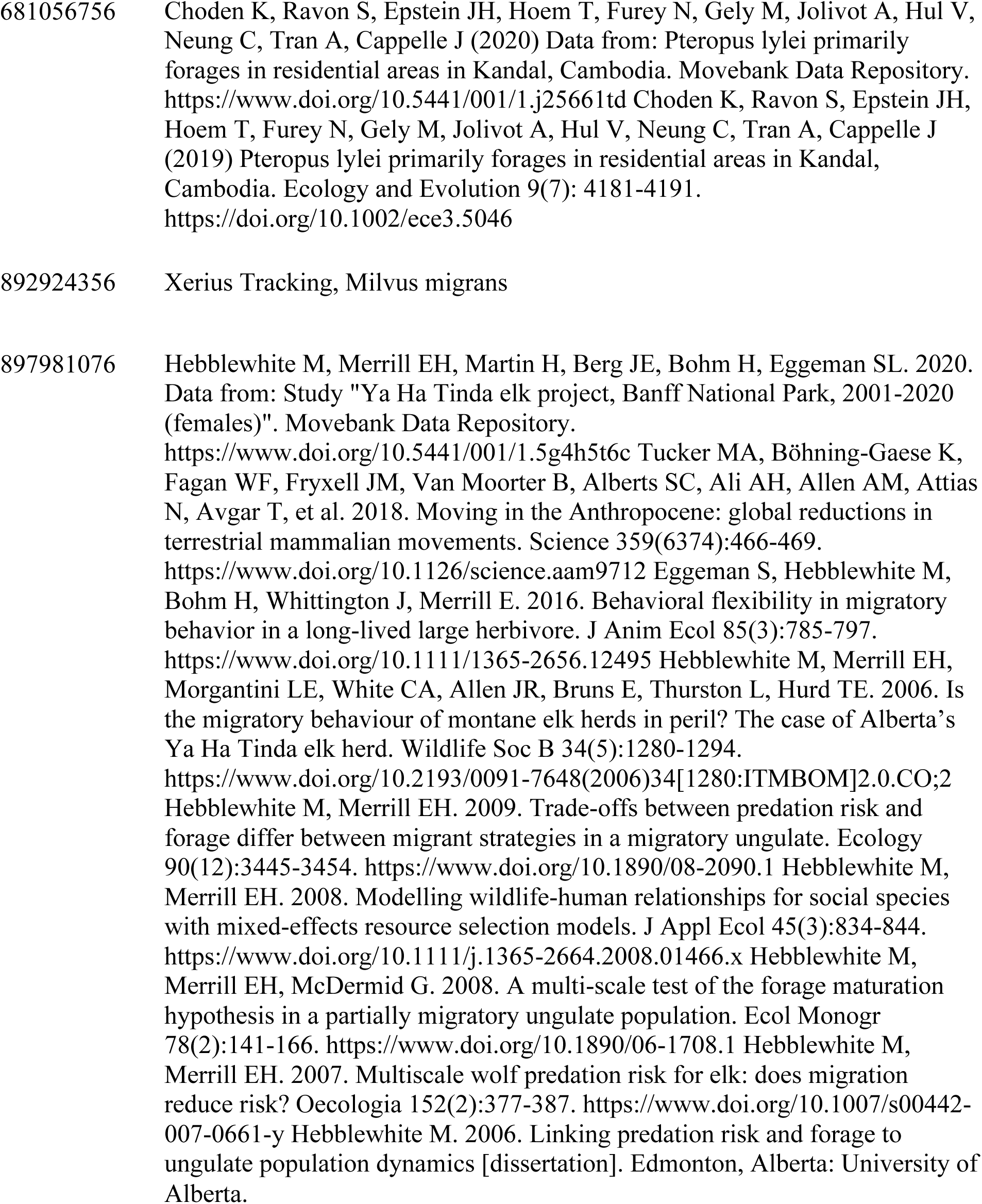
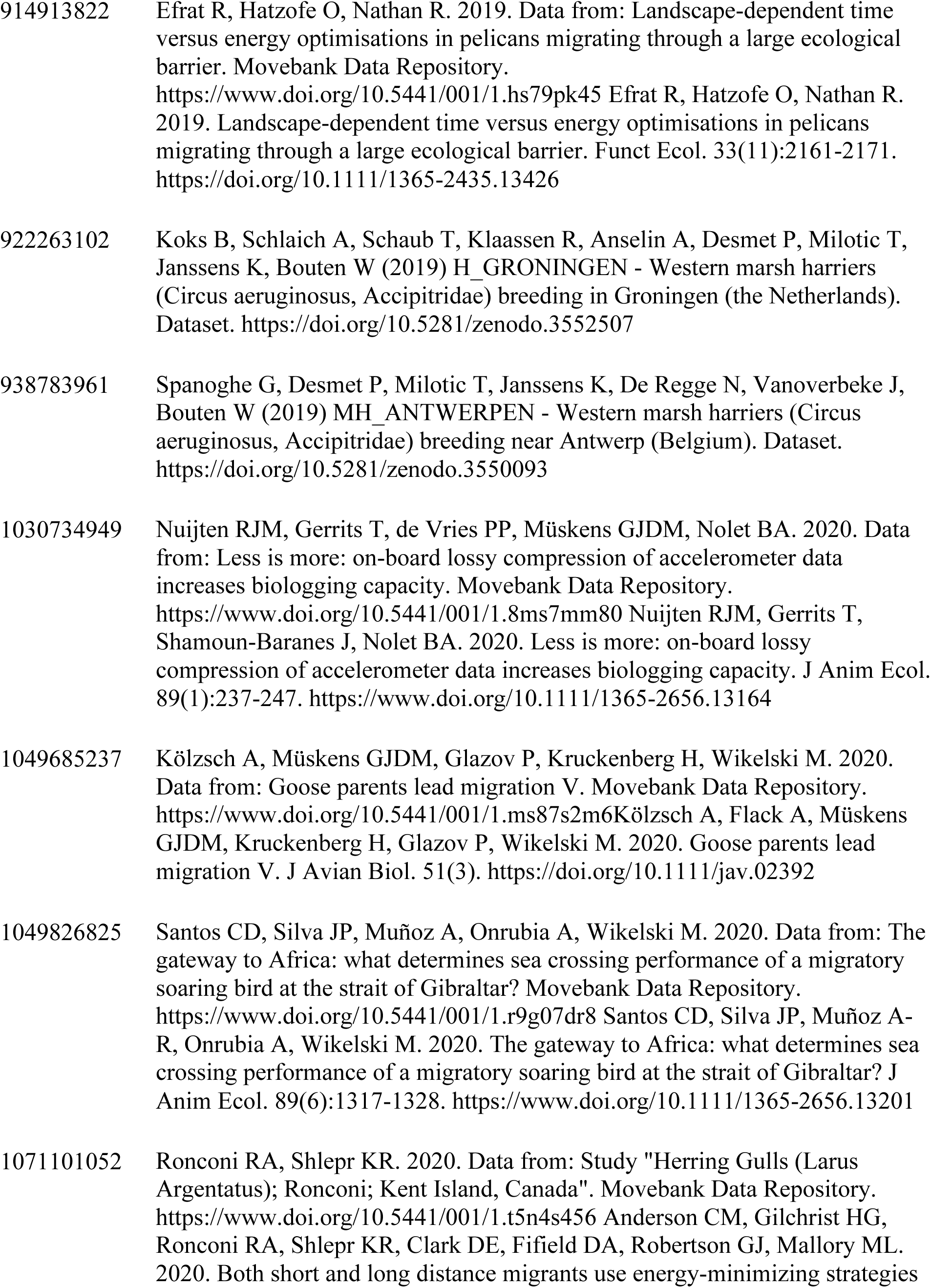
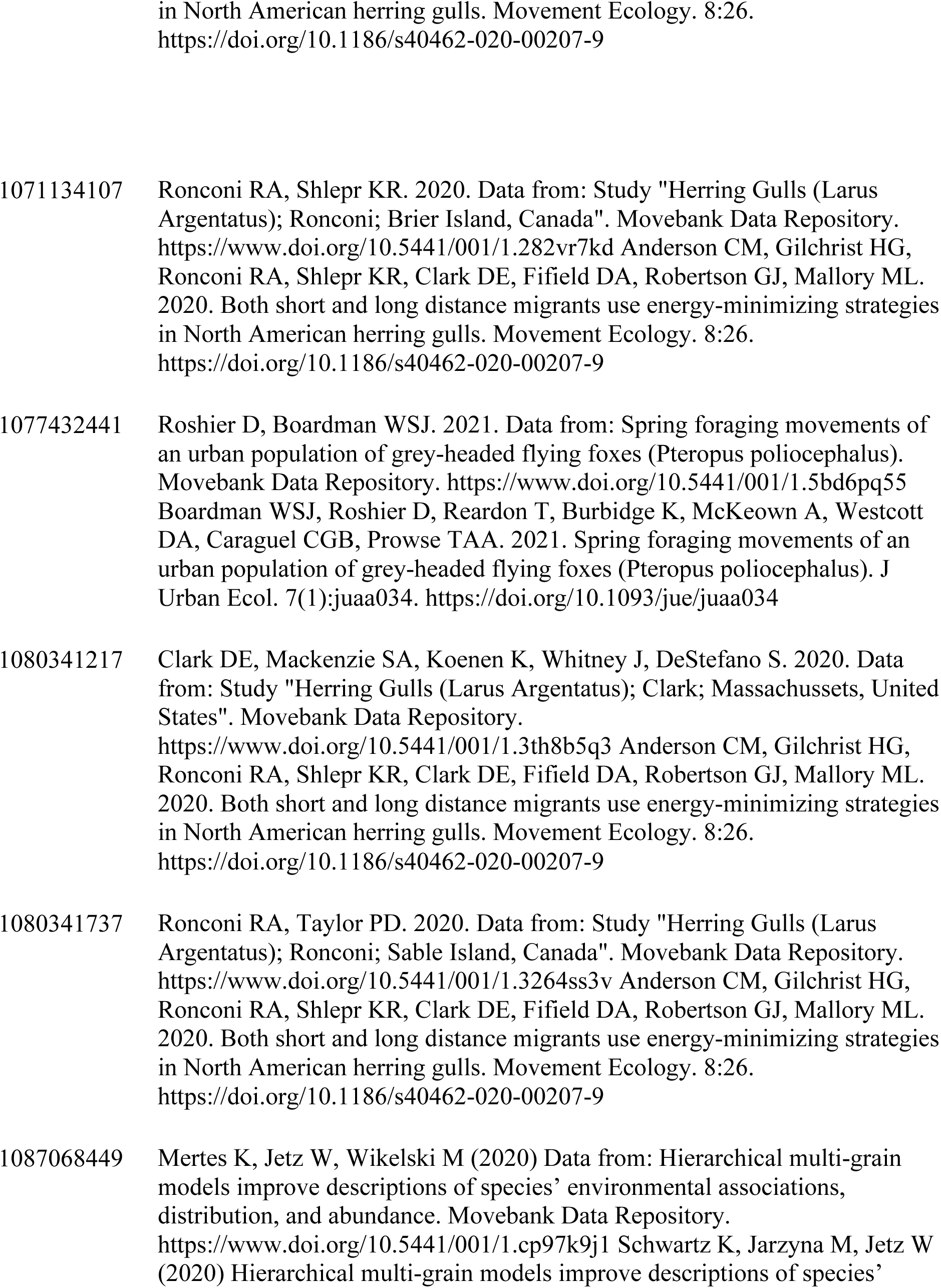
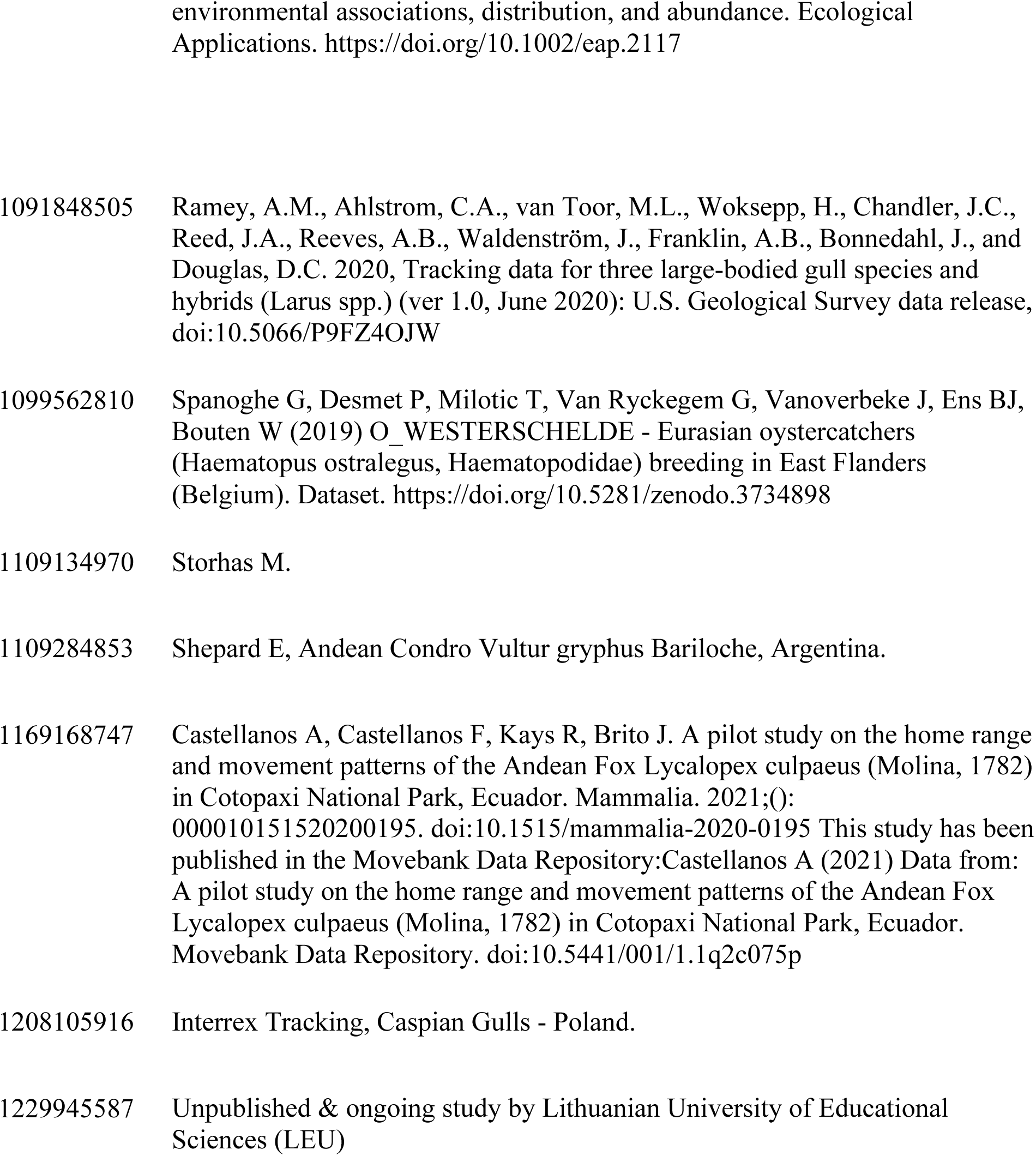
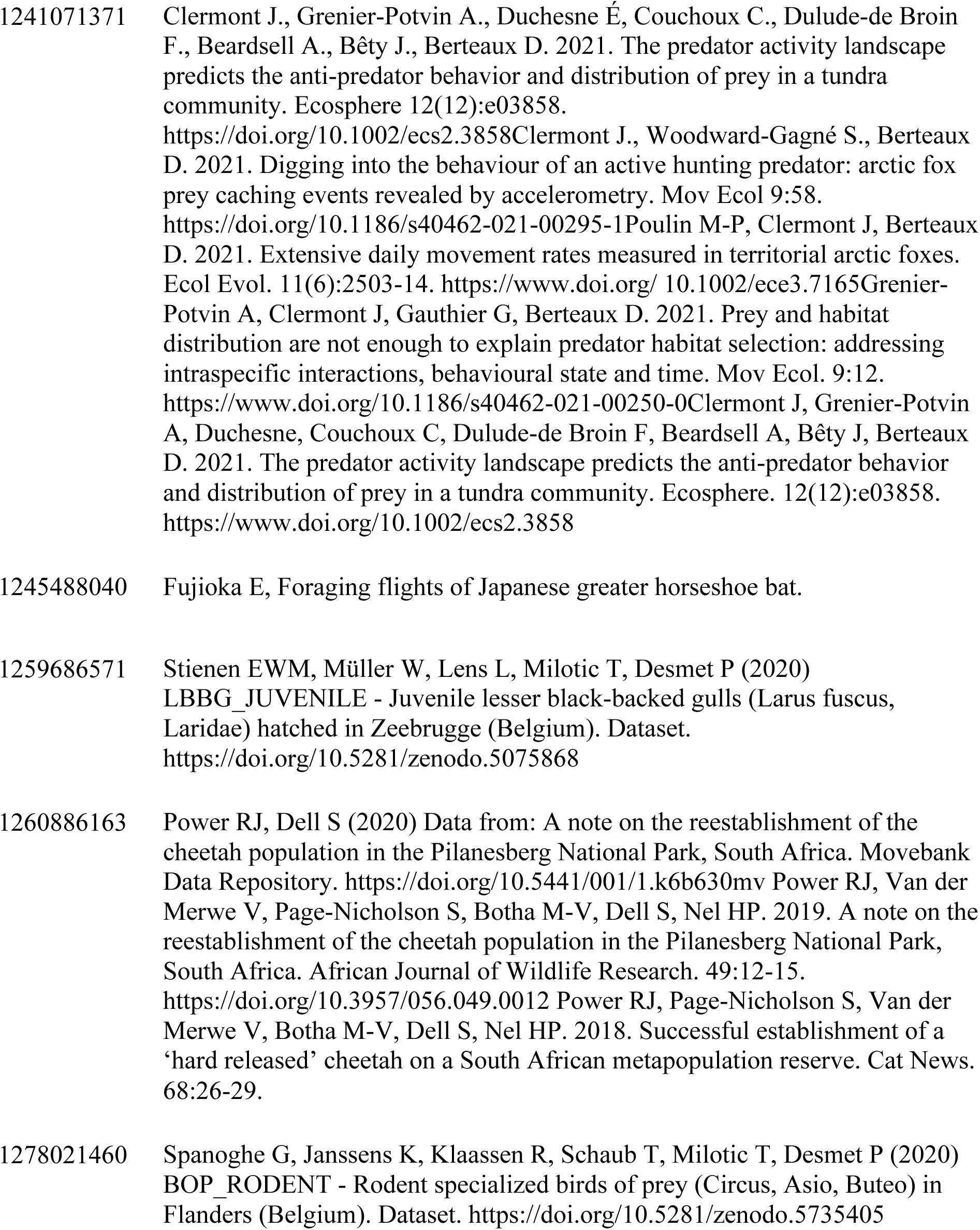
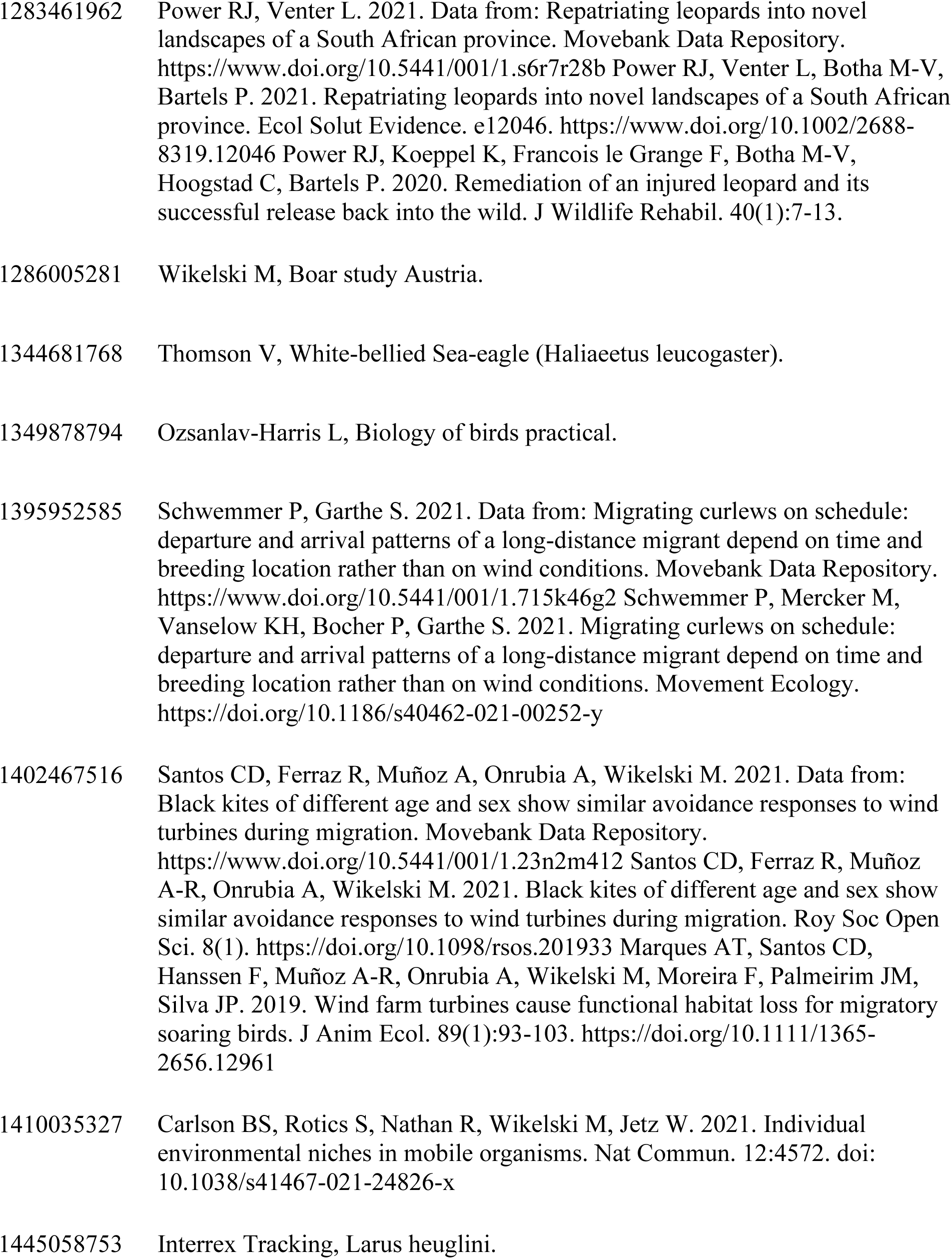
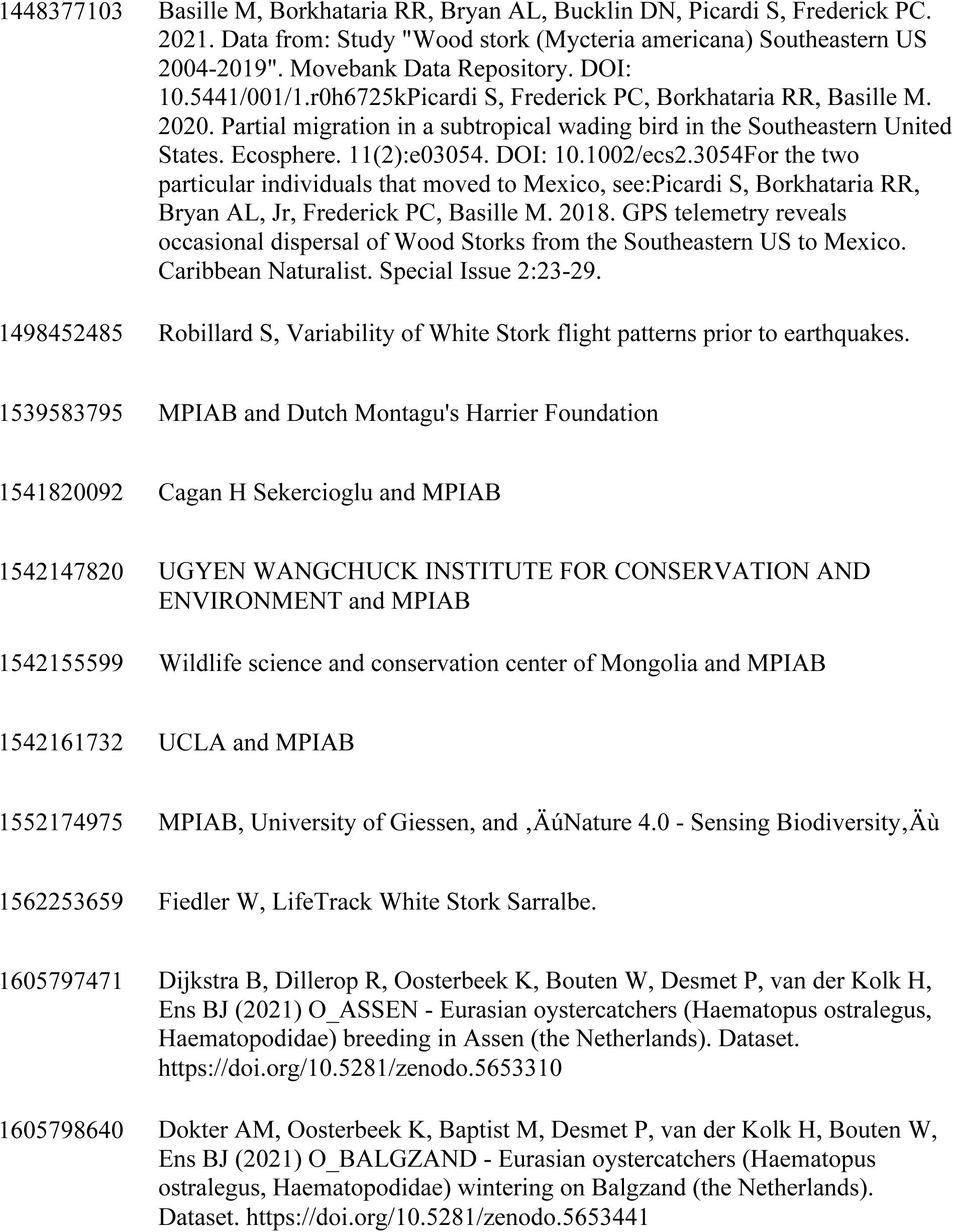
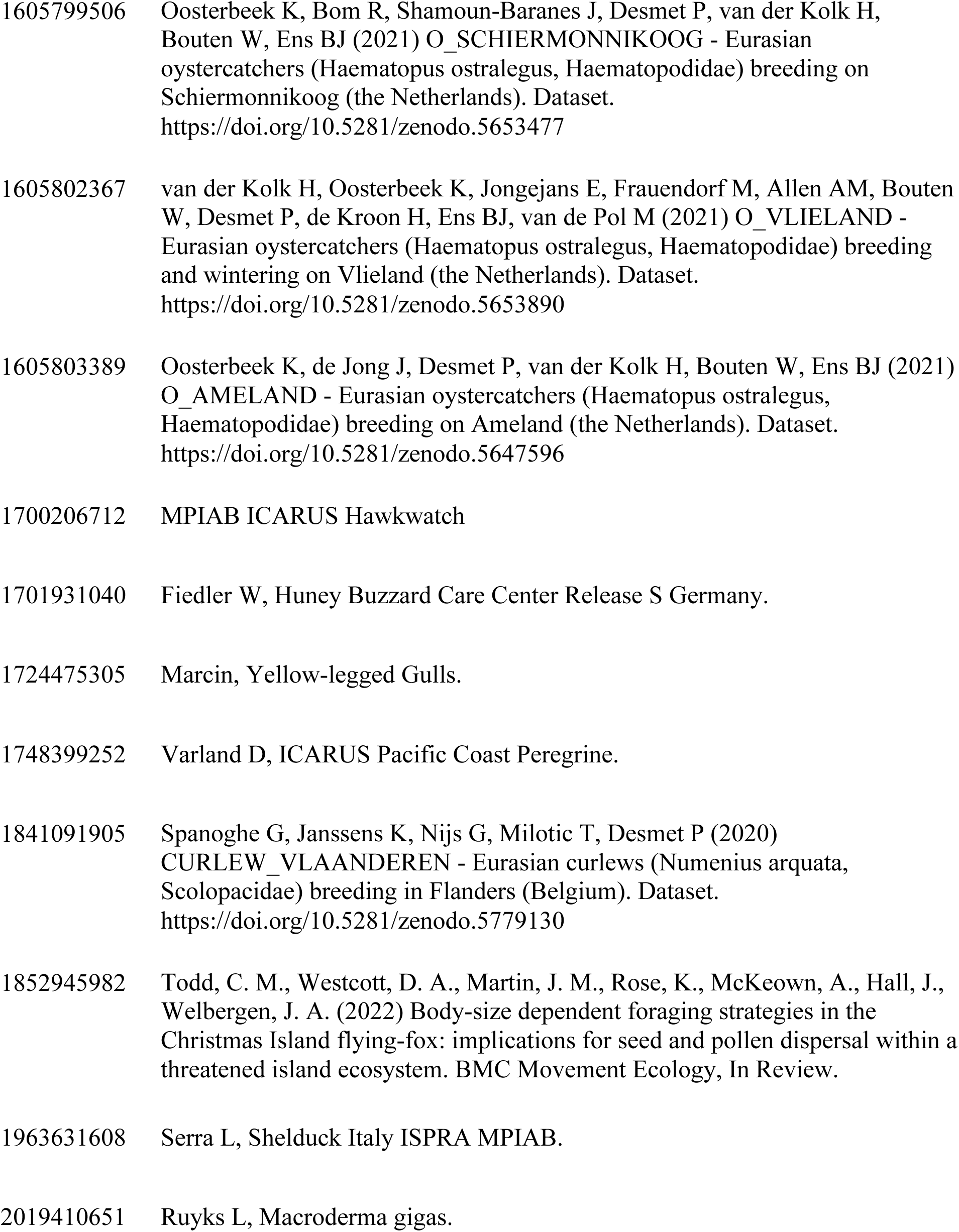
Animal movement datasets accessed via MoveBank under CC0 licenses, including numeric ID and available project names or study references.

